# The microbiome of Antarctica’s endemic chironomid midge: from general microbial community to endosymbiotic bacteria

**DOI:** 10.64898/2025.12.13.694143

**Authors:** Pavlo Kovalenko, Mariia Pavlovska, Yevheniia Prekrasna-Kviatkovska, Anton Puhovkin, Iryna Kozeretska

**Author notes:** Correspondence: Pavlo Kovalenko. Pavlo Kovalenko, Mariia Pavlovska, and Yevheniia Prekrasna-Kviatkovska are contributed equally to this work.

## Abstract

**Background:** The interactions between the host organism and the microbiota play an important role in the host’s survival, influencing its physiology and adaptation. The Antarctic, due to its extreme living conditions, is a unique region for studying such interactions. However, research on the microbiomes of Antarctic terrestrial and freshwater invertebrates remains limited, especially with regard to the region’s sole endemic insect, *Belgica antarctica.* Our aim was to study the *B. antarctica* microbiome collected from different substrates and from varying ornithogenic influence levels.

**Methods:** A total of 330 *B. antarctica* larvae and moss substrates were sampled at three sites with different ornithogenic impact, followed by DNA extraction using Qiagen kits. The V3-V4 region of bacterial 16S rRNA was amplified by PCR and sequenced on the Illumina NovaSeq platform, followed by bioinformatic processing in QIIME2 for OTU clustering, taxonomic classification, and diversity analysis. Further analysis and visualization was performed in R, Python and GraphPad Prism.

**Results:** Amplicon sequencing revealed rich microbial diversity, with NMDS demonstrating clear clustering of samples according to their origin (insects vs substrate) and ornithogenic influence level. Comparative analysis revealed that the vast majority of OTUs associated with *B. antarctica* (ca. 95%) were shared with substrates, whereas the insect-specific fraction was small, highly variable, and partially influenced by moss species. Ornithogenic influence significantly structured the symbiotic microbiota in both insects and substrates, reducing microbial diversity and altering the relative abundance of major bacterial taxa. Overall, substrate type and ornithogenic impact are significant factors influencing microbial community structure, while insect-specific taxa formed only a minor, heterogeneous component of this midge microbiome. Additionally, the *Wolbachia* infection was detected in *B. antarctica* larvae for the first time.

**Conclusion:** The substrate that *B. antarctica* inhabits, as well as ornithogenic influence, play a significant role in the shaping of the species’ microbiome. This Antarctic chironomid also maintains a core microbial community. The detection of *Wolbachia* in *B. antarctica* is the first evidence of this endosymbiont in Antarctica.

## Introduction

It is increasingly acknowledged that the microbiome (the microbial community residing in a healthy host) is essential for its overall health and fitness. Members of the microbiota not only support the host’s overall health but also exist in a state of chronic competition for resources and physical survival in a highly dynamic environment. The permanent potential for infection, along with the rapid evolution of microorganisms, poses a potential threat to the host, which, in turn, develops means of controlling and manipulating its microbiome to limit harm and promote benefits [Kolodny et al., 2020; Wilde et al., 2024]. Interactions with the microbial community, particularly bacteria, may play a significant role in the adaptation of living organisms to challenges during climate change, as the key role of microbiota in host physiology and adaptation has been recognized in recent years [McFall-Ngai et al., 2013; Bang et al., 2018; Haider et al., 2025].

For at least some insect species, interaction with their microbiota is vital for normal development and survival, as associated bacterial microorganisms are involved in processes such as nutrition (e.g., *Acetobacter pomorum*, *Lactobacillus plantarum*, *Serratia symbiotica*, *Staphylococcus gallinarum*, *Ishikawaella*, *Wolbachia*, some Bacteroidetes, Enterobacterales, and Firmicutes representatives, etc.), detoxification (e.g., *Hamiltonella defensa*, *Acetobacter*, *Komagataeibacter*, *Pseudomonas*, and *Sphingomonas*), and protection from a wide range of biotic and abiotic environmental stress factors (e.g., *Buchnera aphidicola*, *Burkholderia gladioli*, *Serratia symbiotica*, *Shikimatogenerans silvanidophilus*, *Hamiltonella*, *Nardonella*, *Regiella*, *Spiroplasma*, *Sphingomonas*, *Wolbachia*, etc.) [Douglas, 2015; Chabanol & Gendrin, 2024]. A large portion of the microbiota associated with a host insect may have facultative, non-obligatory relationships with the host, although both the host and the bacterium can benefit from this association.

Moreover, there are also obligate endosymbiotic bacteria that often play an important role in the adaptation of multicellular organisms [Duron et al., 2008; Massey & Newton, 2022; Serga et al., 2024]. Obligate endosymbionts are able to modify the host phenotype that increases rates of the infection transmission from mother to offspring [Wernegreen, 2012]. Among the most studied are *Wolbachia*, *Cardinium*, *Rickettsia*, *Arsenophonus*, and *Spiroplasma* [Duron et al., 2008; Massey & Newton, 2022]. Representatives of these bacterial genera are also interesting because they are known reproductive parasites in arthropods [Duron et al., 2008]. Although they are found in various arthropod species from different parts of the world, except for the *Rickettsia* in the invasive chironomid midge for Antarctica *Eretmoptera murphyi* Schäffer, 1914 [Brayley et al., 2025a, b], their presence in terrestrial Antarctic ecosystems remains unclear.

Antarctic terrestrial ecosystems are unique due to the sustained biological isolation of Antarctica and the region’s distinctive extreme environmental conditions [Chown et al., 2015; Fraser et al., 2018; Convey & Peck, 2019; Bargagli, 2020; Convey & Biersma, 2024]. The harshness of the Antarctic environment significantly narrows the species diversity of its terrestrial ecosystems, which consist of only a few groups of organisms [Convey et al., 2014; Convey & Biersma, 2024]. In general, the terrestrial Antarctic fauna consists predominantly of invertebrates, with the native fauna comprising Acari (>100 species), Collembola (ca. 25 species), Insecta (2 species), Nematoda (ca. 60 species), Tardigrada (>70 species), and Rotifera (>150 species) [Serga et al., 2024]. Nevertheless the species richness of this group is considerably lower in specific regions, for instance, despite a total of 20 Collembola species being known for the West Antarctic Peninsula region, only 1–3 species are usually recorded on separate islands [Protsenko et al., 2025]. Consequently, studying the microbiome of Antarctic invertebrates and its role in their adaptation to environmental conditions could be of special interest.

Among the two native insects of Antarctica, the sole endemic is the wingless *Belgica antarctica* Jacobs, 1900 [Kozeretska et al., 2022]. This Antarctic chironomid attracts particular interest as a model organism for studying the adaptations of living organisms to a harsh environment [Kozeretska et al., 2022]. This midge is characterized by a relatively long life cycle, lasting two years, with the majority of this period spent in the larval stage, during which midges can feed on a wide range of food resources, including microorganisms, decaying plant material, fungi, and algae [Kozeretska et al., 2022]. Consequently, *B. antarctica* larvae can inhabit various subformations and substrate types, such as algal mats, bryophyte carpets, and seabird nesting areas [Atchley & Davis, 1979; Convey & Block, 1996; Potts et al., 2020; Kovalenko et al., 2021; Kozeretska et al., 2022]. Even though this insect is currently the subject of numerous studies, the research targeting its microbiome and possible host-microbiome interactions is still scarce and significant gaps remain in this area.

Maistrenko et al. (2023) identified 20 bacterial species from five phyla (Actinobacteria, Bacteroidetes, Firmicutes, Fusobacteria, and Proteobacteria) in 34 *B. antarctica* samples. However, the authors noted that only 14 identified species were not potential contaminants from human microbiota, as they are found either in the substrate or in water. Additionally, efforts to identify the endosymbiont *W. pipientis* [Maistrenko et al., 2023; Serga et al., 2024; see also Holmes et al., 2019, discussion section] associated with *B. antarctica* have yielded weak or negative results. Based on this information, the composition of the *B. antarctica* microbiome remains a subject of debate. It is unclear whether the microbiome is primarily dependent on microorganisms from the surrounding substrate, which also serves as the insect’s trophic supply, or if a more species-dependent core microbiome exists.

The role of the microbiome is also of significant interest given the environmental changes that Antarctic ecosystems are currently undergoing, specifically climate change [Turner et al., 2014; Convey & Peck, 2019; Siegert et al., 2019; Bargagli, 2020]. Among the indigenous Antarctic species expanding into new territories due to climate changes, the gentoo penguin *Pygoscelis papua* (J.R.Forster, 1781) stands out. This species has significantly increased its population and expanded its range southward over the past 20 years [Lynch et al., 2012; Korczak-Abshire et al., 2021], a trend also observed at Galindez Island [Parnikoza et al., 2018]. This ornithogenic impact from penguins not only alters the chemical composition of the substrate but also changes the taxonomic composition of microbial communities within moss banks [Prekrasna-Kviatkovska et al., 2024]. The implications of these shifts for the microbiome of invertebrates inhabiting the transformed ecosystems remain an open question.

The objective of this study was to investigate the microbiome of *B. antarctica* collected from different substrates and subjected to varying intensities of ornithogenic impact. We aimed to assess the degree to which the insect’s microbiome is influenced by the substrate, ornithogenic impact, and to identify the potential infection of *B. antarctica* with the most well-known obligate endosymbionts of arthropods (*Wolbachia*, *Rickettsia*, *Cardinium*, *Spiroplasma*, and *Arsenophonus*).

## 2. Materials and Methods

### 2.1. Samples collection and fixation

Sampling was conducted during March-April 2023 at the three sites on Galindez Island (Argentine Islands, Wilhelm Archipelago) (Fig. 1), characterized by varying levels of ornithogenic influence: the first site (65.244736 S, 64.256188 W), where a colony of penguins *P. papua* has been located since 2007 – high ornithogenic influence; the second site (65.246031 S, 64.248037 W) was next to a new colony of *P. papua* that has been located since 2019 – medium ornithogenic influence; the third site (65.248946 S, 64.246873 W) – a location without bird colonies, with only a nesting territory of a south polar skuas *Stercorarius maccormicki* H.Saunders, 1893 pair nearby – low ornithogenic influence (Fig. S1).

**Figure 1.**
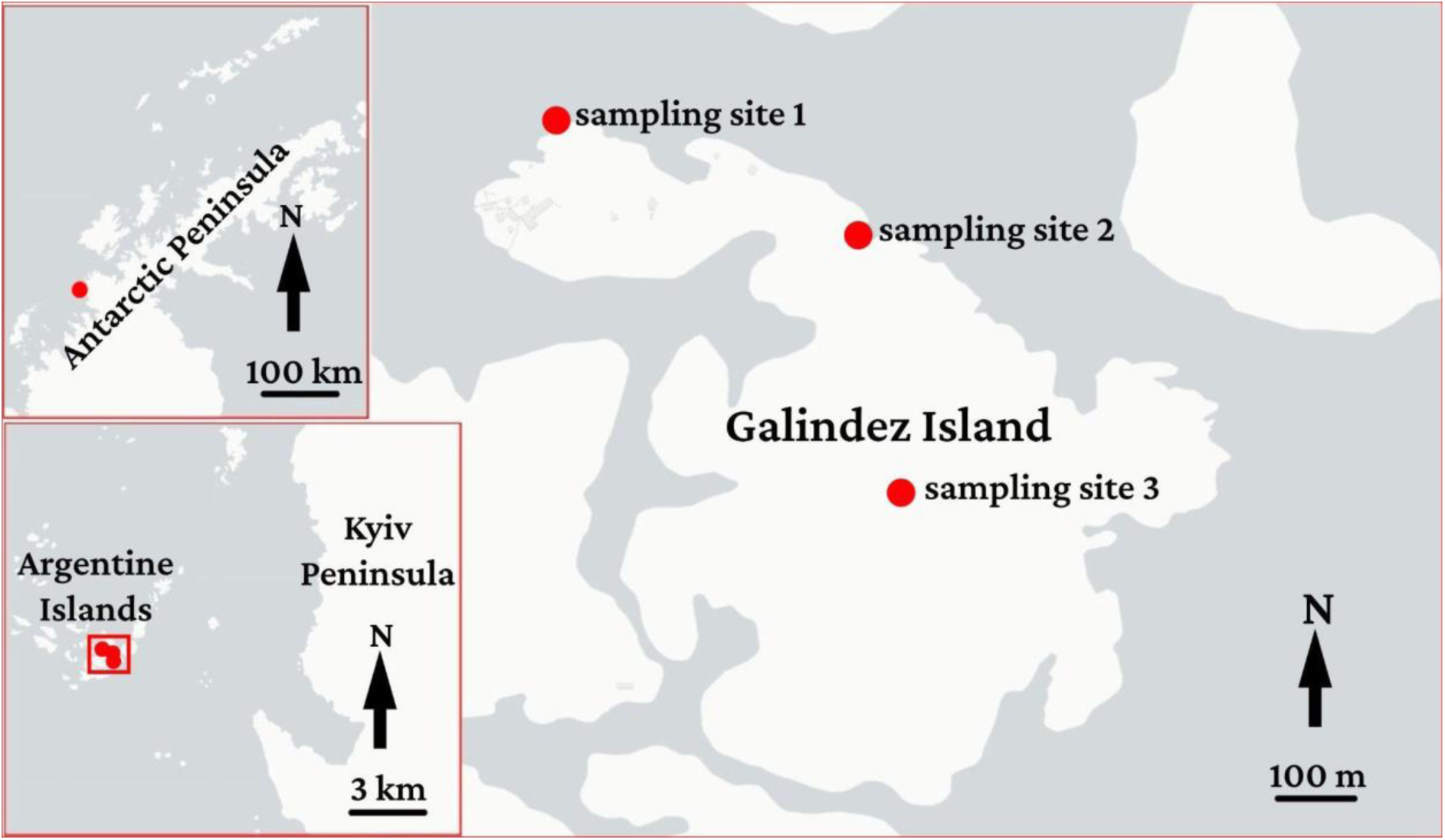
Map indicating the sampling sites

From each location, two types of moss substrate were collected: 1) *Sanionia georgicouncinata* (Müll. Hal.) Ochyra & Hedenäs, 1998 and 2) *Warnstorfia fontinaliopsis* (Müll. Hal.) Ochyra, 2001, with at least three replicates (ten specimens each) of *B. antarctica* larvae taken from every substrate (33 replicates total). The substrate was preserved in 15 ml Falcon-type tubes and stored in DNA/RNA Shield solution, while the insect larvae were fixed in 96% ethanol. Additionally, nine replicates of *B. antarctica* larvae samples collected from *W. fontinaliopsis* substrate at the sampling site 1 and both studied substrates at the sampling site 3 were fixed in DNA/RNA Shield solution, as the substrate (Table 1). All samples were refrigerated at +4°C for 12 hours after the fixation, frozen at -20°C for storage, and then transported to Ukraine for further analysis.

**Table 1.**
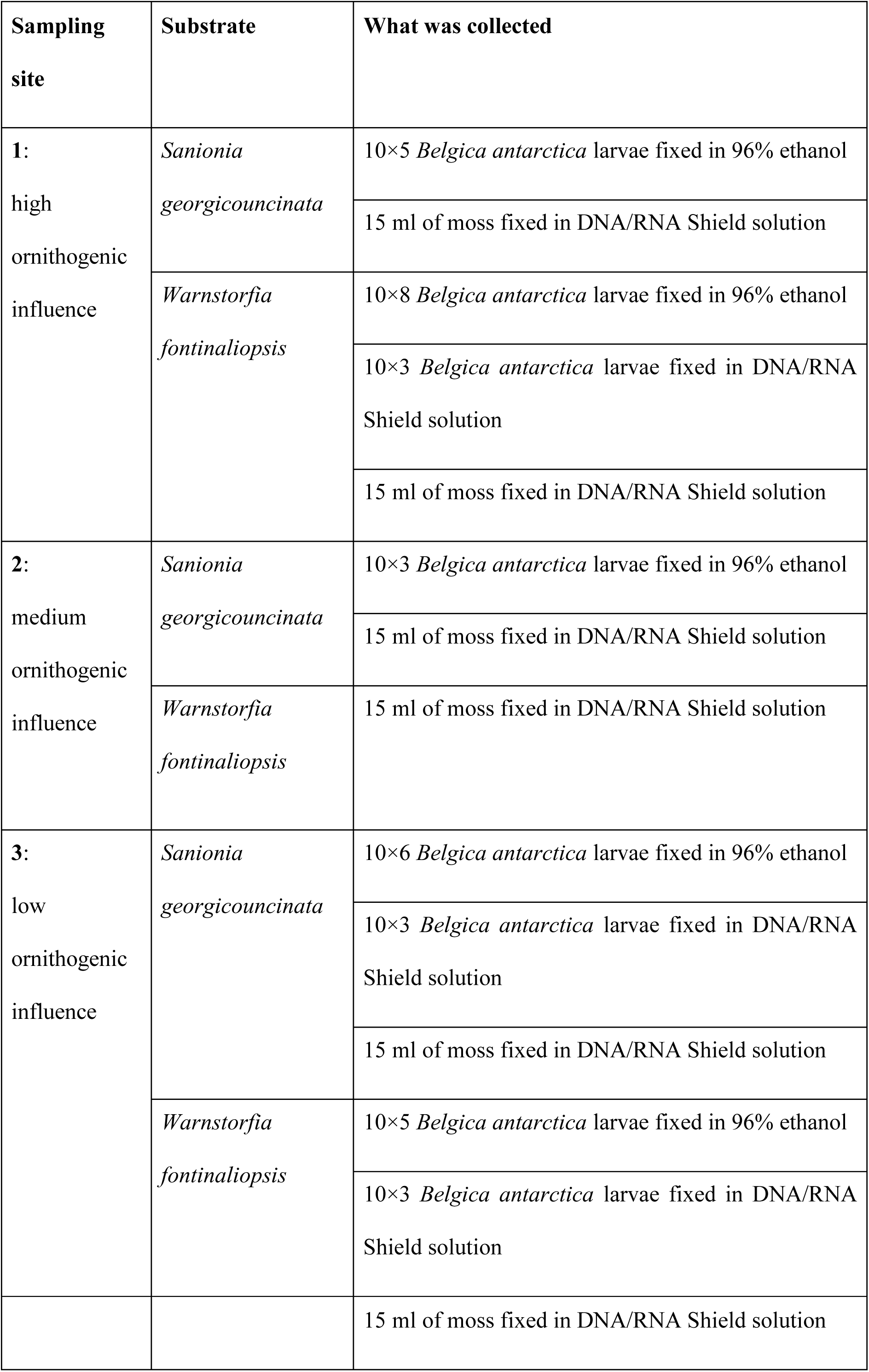
Overview of sampling across sites with varying ornithogenic impact, substrate and storage solution.

| Sampling site | Substrate | What was collected |
| --- | --- | --- |
| 1:<br>high<br>ornithogenic<br>influence | *Sanionia georgicouncinata* | 10×5 *Belgica antarctica* larvae fixed in 96% ethanol |
|  |  | 15 ml of moss fixed in DNA/RNA Shield solution |
|  | *Warnstorfia fontinaliopsis* | 10×8 *Belgica antarctica* larvae fixed in 96% ethanol |
|  |  | 10×3 *Belgica antarctica* larvae fixed in DNA/RNA Shield solution |
|  |  | 15 ml of moss fixed in DNA/RNA Shield solution |
| 2:<br>medium<br>ornithogenic<br>influence | *Sanionia georgicouncinata* | 10×3 *Belgica antarctica* larvae fixed in 96% ethanol |
|  |  | 15 ml of moss fixed in DNA/RNA Shield solution |
|  | *Warnstorfia fontinaliopsis* | 15 ml of moss fixed in DNA/RNA Shield solution |
| 3:<br>low<br>ornithogenic<br>influence | *Sanionia georgicouncinata* | 10×6 *Belgica antarctica* larvae fixed in 96% ethanol |
|  |  | 10×3 *Belgica antarctica* larvae fixed in DNA/RNA Shield solution |
|  |  | 15 ml of moss fixed in DNA/RNA Shield solution |
|  | *Warnstorfia fontinaliopsis* | 10×5 *Belgica antarctica* larvae fixed in 96% ethanol |
|  |  | 10×3 *Belgica antarctica* larvae fixed in DNA/RNA Shield solution |

|  |  |
| --- | --- |
|  | 15 ml of moss fixed in DNA/RNA Shield solution |

### 2.2. DNA extraction

DNA extraction from *B. antarctica* larvae was carried out using the Qiagen DNeasy Blood & Tissue Kit (Qiagen, Germany). Each replicate, consisting of ten individuals (crushed using sterile pestles), was processed as a separate extraction. DNA extraction from plant substrate was performed using the Qiagen DNeasy PowerSoil Pro Kit (Qiagen, Germany). In both cases, the manufacturer’s instructions were followed.

### 2.3. Microbiome analysis

V3-V4 region of bacterial 16S rRNA was sequenced at Novogen (Novogene Co., Ltd., UK) using the Illumina NovaSeq PE250 platform with 515F-806R barcode universal primers [Caporaso et al., 2011; Nowak et al., 2025]. Phusion High-Fidelity PCR Mix (New England Biolabs) was used for the PCR amplification under the following conditions: an initial denaturation at 98°С for 1 min, followed by 30 cycles of denaturation at 98°С for 10 s, annealing at 50°С for 30 s, and elongation at 72°С for 60 s, with a final elongation step at 72°С for 5 min. The resulting PCR products were analyzed on a 2% agarose gel and purified using the Qiagen Gel Extraction Kit (Qiagen, Germany). Illumina NEBNext Ultra DNA Library Preparation Kit was subsequently used for library preparation, followed by library quality assessment with the Qubit 2.0 fluorometer (Thermo Scientific, USA) and Agilent Bioanalyzer 2100 (Agilent, USA) system. Sequencing data generated for the current study have been deposited in the NCBI database under accession number PRJNA1374202.

### 2.4. Bioinformatic and statistical analysis

Sequences were merged and oriented in the 5’-3’ direction using FLASH (v1.2.7) [Magoč & Salzberg, 2011]. Sequence processing was carried out in QIIME2 2019.7 [Bolyen et al., 2019], where barcodes, homopolymers, chimeric reads, and short sequences (<150 bp) were removed. Operational taxonomic units (OTUs) were identified through de novo clustering at 97% similarity using the UPARSE algorithm (Uparse v7.0.1090) [Edgar, 2013]. Taxonomic classification of OTUs was performed using the Silva SSU Ref NR 99 database (v.138) [Quast et al., 2013].

The taxonomic structure of the *B. antarctica* microbiome was analyzed under the differential influence of ornithogenic factor, substrate characteristics and storage solution type.

Alpha diversity metrics, including the Shannon diversity index and phylogenetic distance (PD), were calculated using the “Diversity core metrics” plugin in QIIME2 2019.7. The Shapiro-Wilk test [Shapiro & Wilk, 1965] in R (v.2.15.3) (R Core Development Team) was used to assess data normality, followed by Kruskal-Wallis test [Kruskal & Wallis, 1952] to detect significant differences in diversity of the samples. A non-metric multidimensional scaling (NMDS) plot with Bray-Curtis measure [Legendre & Legendre, 2012] was used to visualize microbial community differentiation across substrates, while bacterial taxa with the highest contribution to the detected difference were depicted with principal component analysis (PCA) [Jolliffe, 2002]. Mantel test with Spearman correlation [Legendre & Legendre, 2012] was performed to disentangle the impact of substrate type and sample preservation method on microbial community taxonomic structure. The contribution of environmental parameters to the observed microbial community differentiation was visualized with the constrained ordination (CAP) using the *vegan* package in R [Oksanen et al., 2025].

The taxonomic structure and differentiation of microbial communities was visualized in GraphPad Prism (v10.0.0, GraphPad Software, Boston, Massachusetts, USA).

## 3. Results

### 3.1. Diversity of *Belgica antarctica* microbiomes in context of ecologically different environment

Amplicon sequencing resulted in an average of 168330.33 qualified reads per sample with the minimum of 56383 and the maximum of 214954 (Table S1). The rarefaction curves showed that the sequencing was deep enough to capture the overall microbial diversity (Figs. S2A, S1B).

NMDS on OTUs dataset revealed the presence of separate clusters corresponding to the origin of the sample (insect vs. moss) and ornithogenic impact on the substrate (high, medium or low) (Fig. S3).

Consequently the dataset was split into the following categories in order to detect the impact of substrate on *B. antarctica* microbiota: 1) the OTUs shared between moss and insect (Insect and Warnstorfia Mutual; Insect and Sanionia Mutual), referred hereinafter as Mutual microbiota; 2) the OTUs unique for the insect (Insect and Warnstorfia Unique; Insect and Sanionia Unique) – Unique Insect and 3) the OTU shared between the insects from different substrates (Insects Mutual) – Mutual Insect (Fig. 2A).

**Figure 2.**
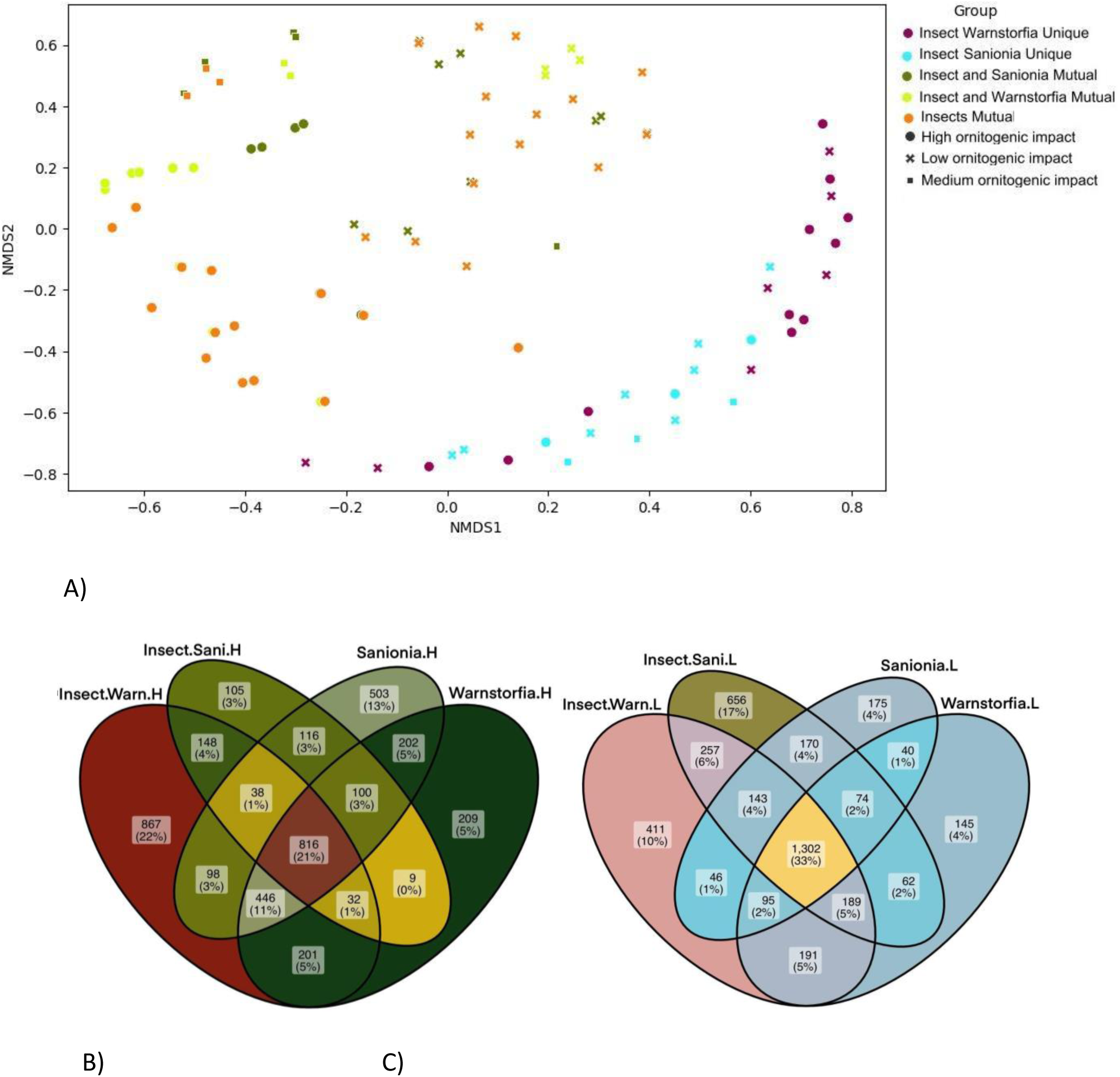
A) Clustering of different samples’ groups based on the NMDS analysis (squares – medium, crosses – high and circles – low ornithogenic impact). Insect Warnstorfia Unique – the OTUs unique for *B. antarctica* from the *Warnstorfia fontinaliopsis*; Insect Sanionia Unique – the OTUs unique for *B. antarctica* from the *Sanionia georgicouncinata*; Insect and Sanionia Mutual – the OTUs shared between *B. antarctica* from the *Sanionia georgicouncinata* and its substrate; Insect and Warnstorfia Mutual – the OTUs shared between *B. antarctica* from the *Warnstorfia fontinaliopsis* and its substrate; Insects Mutual – the OTUs shared between *B. antarctica* from different substrates. B) Venn diagram representing the number of OTUs shared between *B. antarctica* and substrate under high ornithogenic impact. C) Venn diagram representing the number of OTUs shared between *B. antarctica* and substrate under low ornithogenic impact: InsectWarn – *Belgica antarctica* the *Warnstorfia fontinaliopsis*, InsectSani – *Belgica antarctica* from the *Sanionia georgicouncinata*

The NMDS on newly-defined groups uncovered the presence of 2 clusters in *B. antarctica* microbial communities – formed by the unique insect OTUs and the OTUs, which are mutual to the insect and to the substrate (Fig. 2A). Notably, high variation was detected between the samples in the Unique Insect group, which represented only 1.7 ± 3.2% of all OTUs, whereas the Mutual groups comprised 95 ± 8.3% on average.

Additionally, NMDS revealed the variation between the Unique Insect OTUs depending on the moss species they originate from, which signals the potential impact of feeding substrate on *B. antarctica’s* own microbiota (Fig. 2A). Yet, no influence of moss species was detected on clustering of Mutual microbiota. Meanwhile, ornithogenic impact was only significant for clustering of Mutual microbiota, having no impact on structuring the unique insect microbiome.

The OTUs shared between the insects from different substrates clustered together with the Mutual microbial communities, being similarly influenced by ornithogenic factor (Fig. 2A).

A total of 1777 OTUs (45%) were Mutual for Insects and *W. fontinaliopsis*, and 1689 OTUs (43%) were Mutual for Insects and *S. georgicouncinata* (Fig. 2C), all collected under low ornithogenic impact. In contrast, when high-impacted samples were analyzed, the numbers decreased to 1495 (38%) and 1070 (29%) OTUs, respectively (Fig. 2B). The highest number of unique OTUs was detected in insects associated with *S. georgicouncinata* under low ornithogenic impact (656 OTUs), and with *W. fontinaliopsis* under high impact (867 OTUs) (Fig. 2B, C).

Kruskal-Wallis test revealed the presence of statistically significant differences in both Shannon (p < 0.001) and Phylogenetic Diversity (p = 0.03) indices between microbial communities originating from insect and moss samples collected under the differential ornithogenic impact (Figs. 3A, B). However, no pairwise comparisons remained statistically significant after Dunn’s post-hoc correction, likely due to the limited sampling size in each group. The lowest average diversity indices were observed in microbial communities of *B. antarctica*, inhabiting in *W. fontinaliopsis* – 4.39 (Shannon) and 58.59 (Faith PD). The diversity metrics for both insects’ and substrate microbial communities dropped under higher ornithogenic impact. In particular, the average for Shannon index was 3.81/4.1 and for PD – 81.1/104.6 in *B. antarctica* microbiome from *W. fontinaliopsis* and *S. georgicouncinata* respectively (Fig. 3).

**Figure 3.**
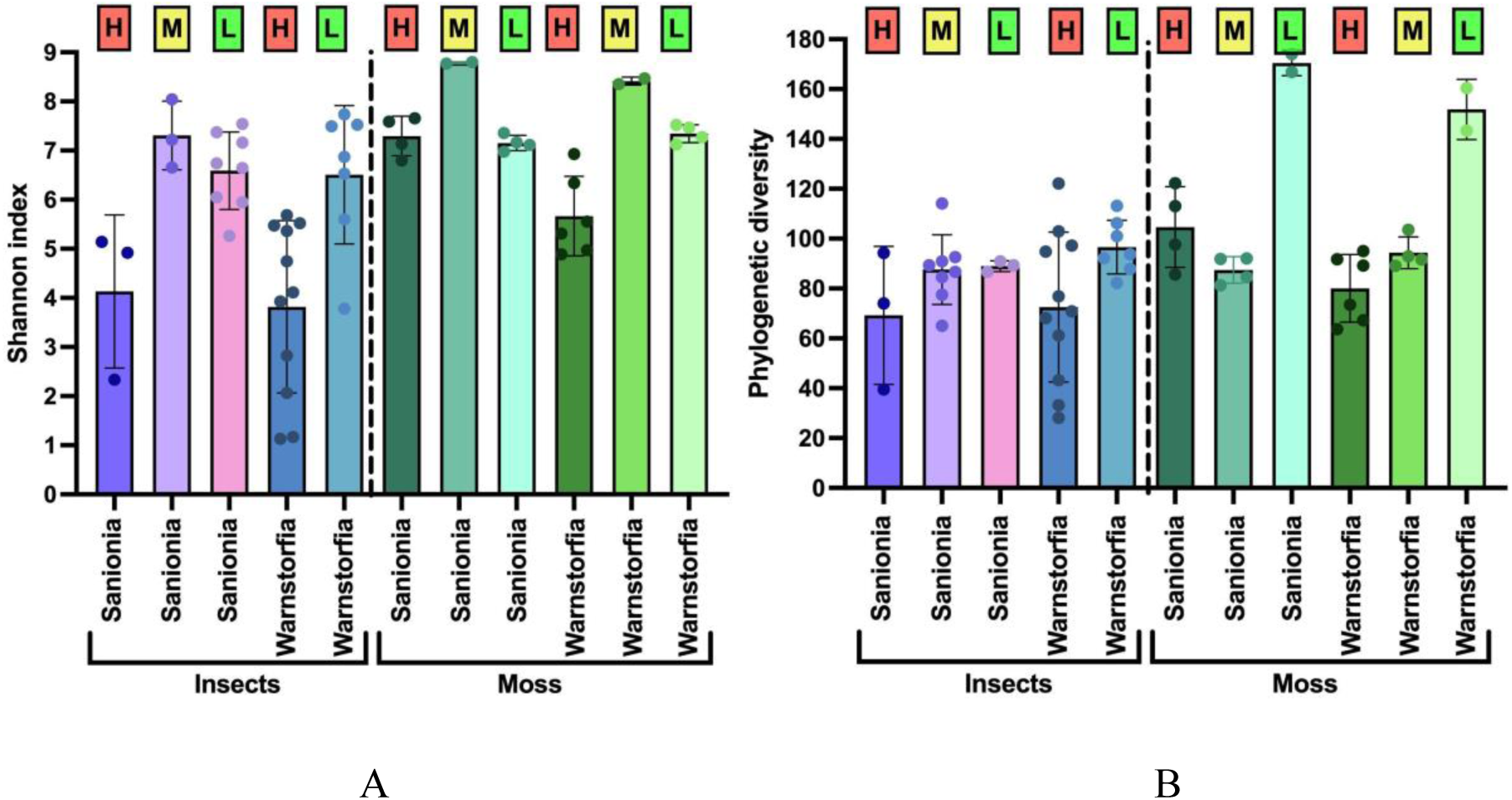
Community diversity indices in different groups of samples: **A**) Shannon index, **B**) phylogenetic diversity (PD). Capital letters above columns indicate the level of ornithogenic impact: L – low, M – medium, H – high

### 3.2. Microbial communities taxonomic structure

We analyzed the taxonomic composition of the microbial communities of insects and substrates, where they were collected (*S. georgicouncinata* and *W. fontinaliopsis*). The taxonomic composition of the microbial communities of *B. antarctica* was shaped by substrate type and ornithogenic influence. Similarly, the microbiomes of the substrates were affected by the ornithogenic factor (Figs. 4, 5, 6, 7).

**Figure 4.**
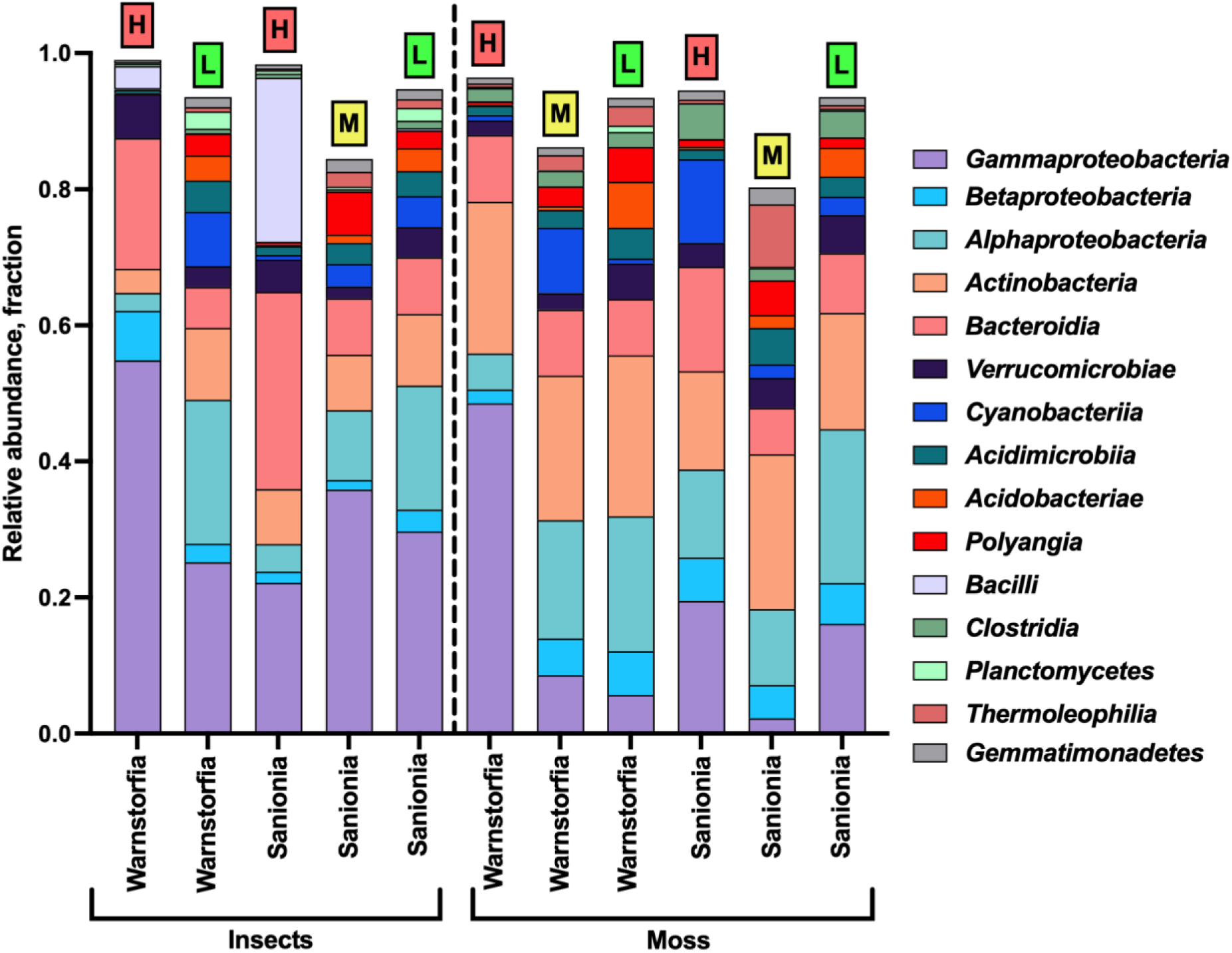
Taxonomic distribution (fraction of the total) of microbial communities at the class level (top 15 most abundant taxa) in different types of samples. Capital letters above columns indicate the level of ornithogenic impact: L – low, M – medium, H – high

**Figure 5.**
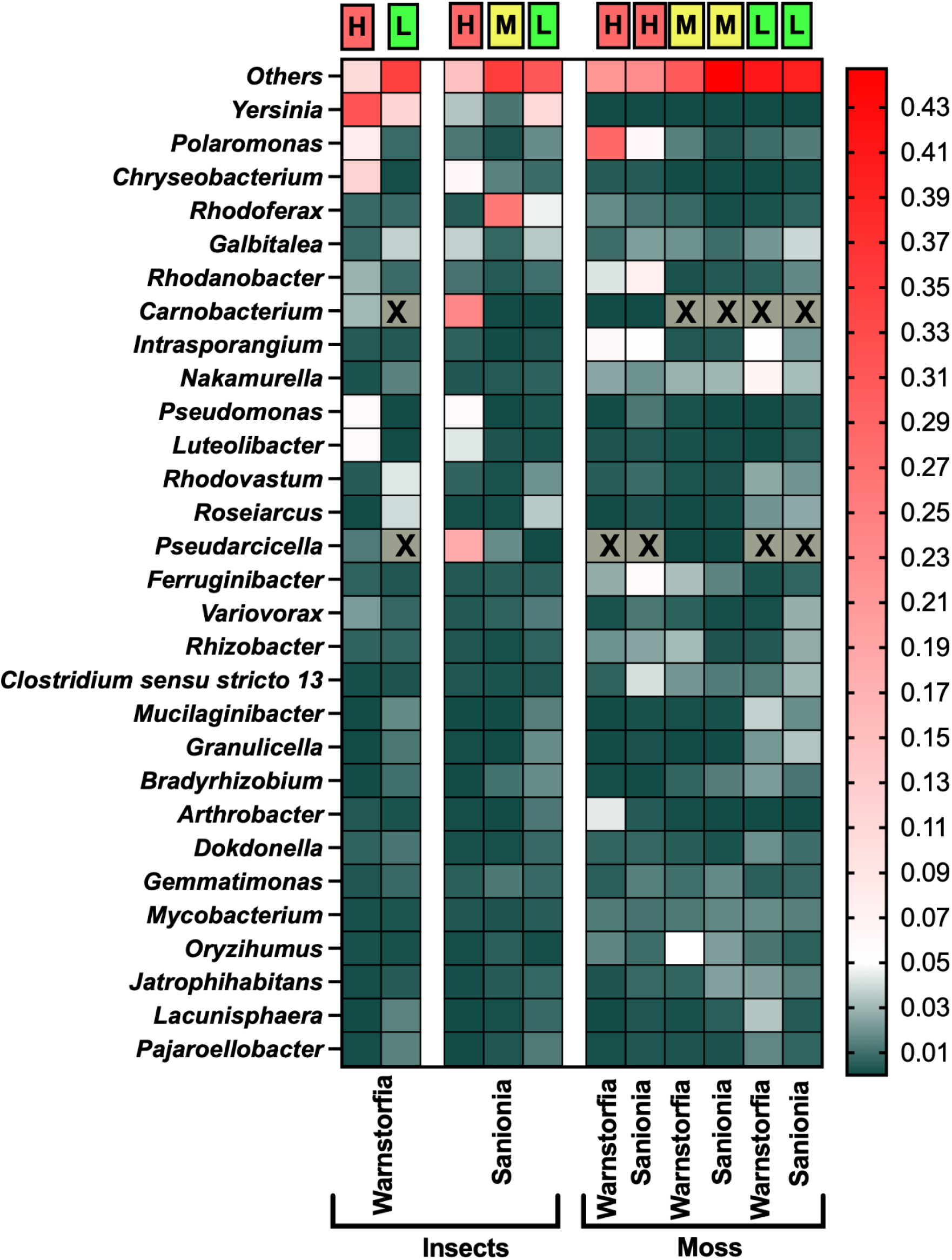
Taxonomic distribution of microbial communities at the genus level (top 30 most abundant genera) in different sample types. Capital letters above columns indicate the level of ornithogenic impact: L – low, M – medium, H – high. Crosses indicate absence of taxa in specific samples

**Figure 6.**
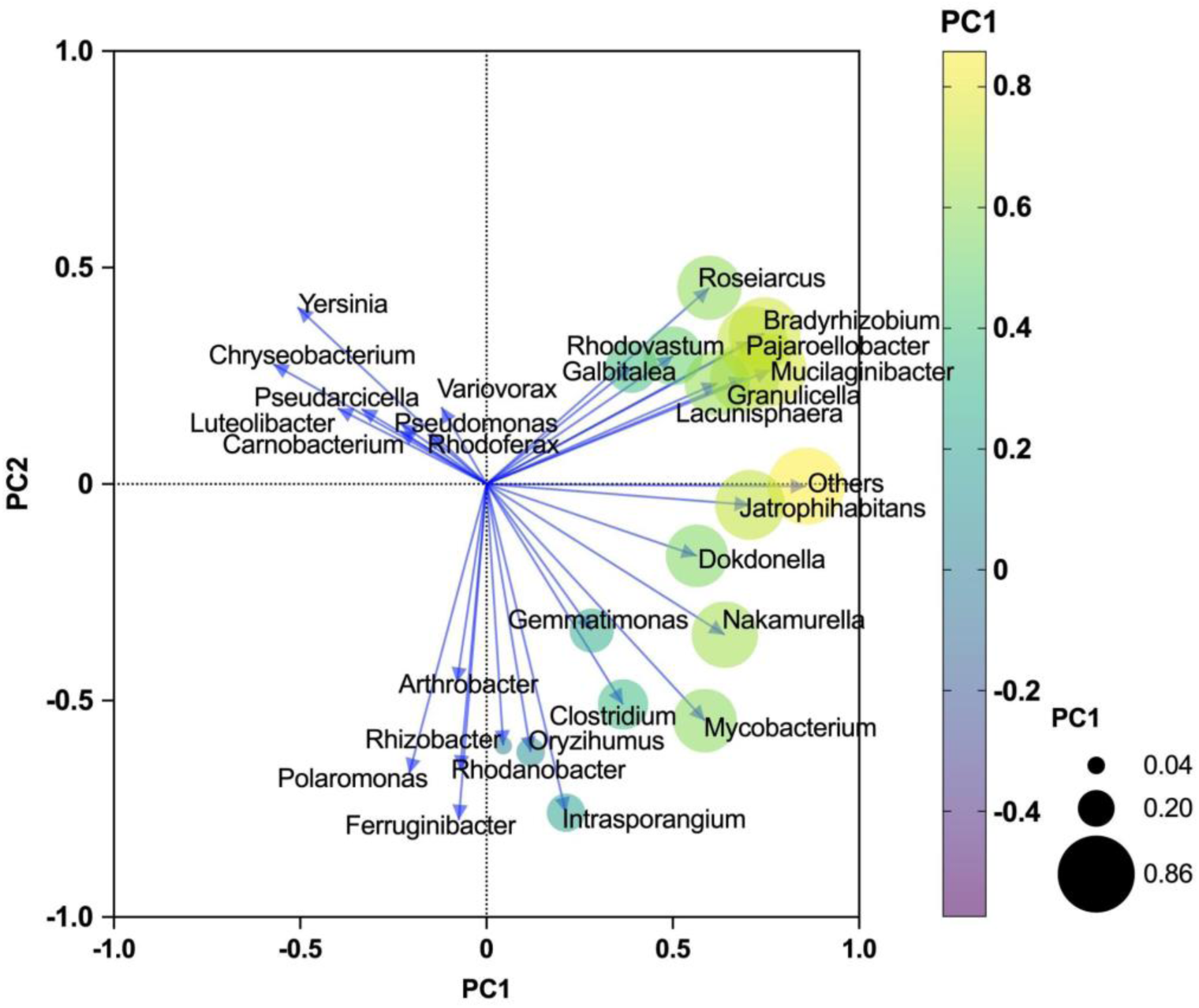
The contribution of microbial genera to the observed community differentiation identified by the PCA

**Figure 7.**
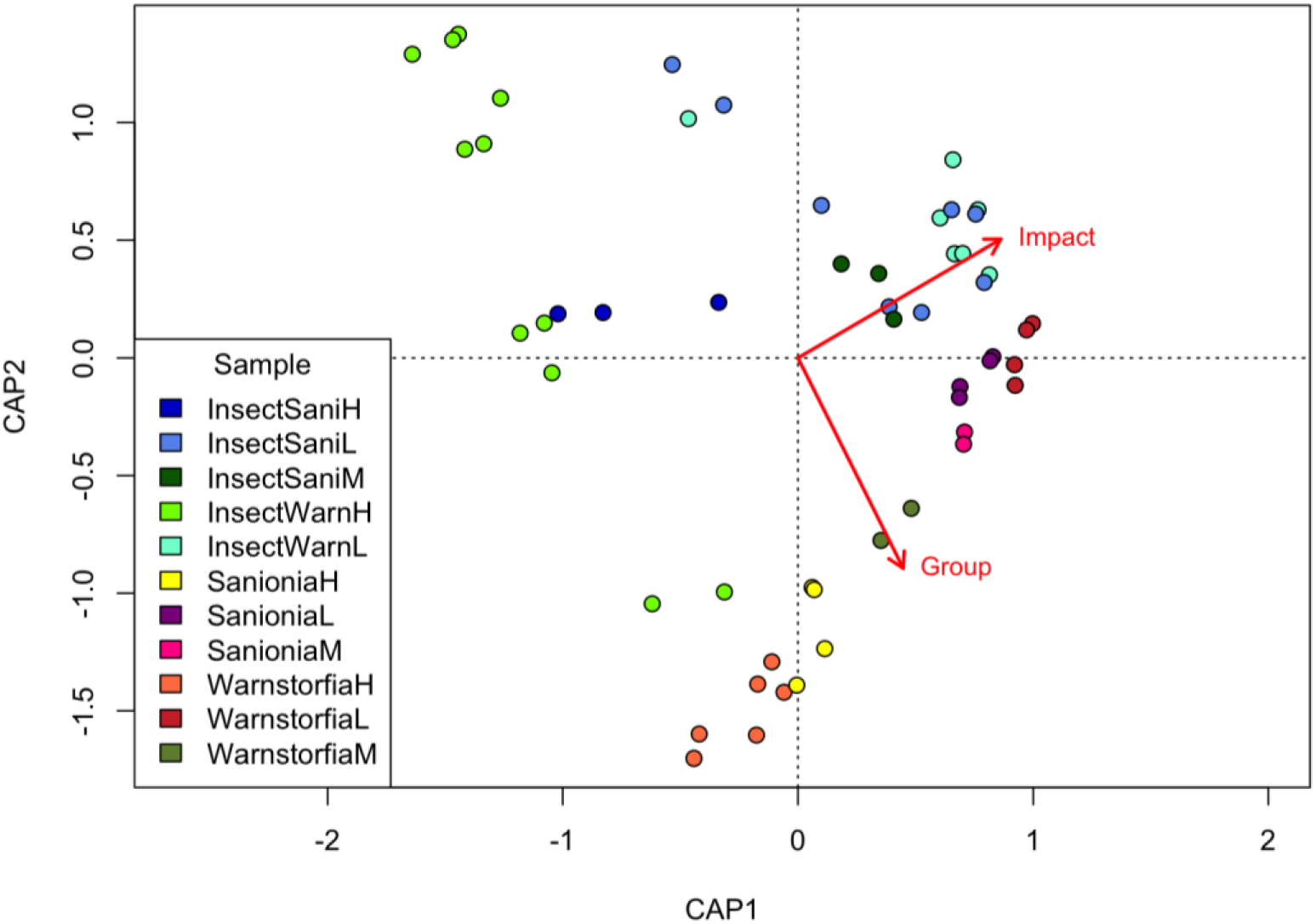
CAP analysis showing the contribution of primary environmental factors to microbial community differentiation: **Impact** – ornithogenic impact: high (H), medium (M), and low (L) and **Group** – microbial communities inhabiting *B. antarctica* or moss. InsectSani and InsectWarn refer to *B. antarctica* larvae collected from *Sanionia georgicouncinata* and *Warnstorfia fontinaliopsis*, respectively; *Sanionia* and *Warnstorfia* – moss genus names representing different substrate types used in the analysis

At the phylum level, Proteobacteria represented the majority in all groups fluctuating in their relative abundance from 38% in the *S. georgicouncinata* substrate to 52% in *B. antarctica* microbiome from *W. fontinaliopsis* samples (Fig. S4). Bacteroidota were second most abundant in the microbiome of insects collected in *W. fontinaliopsis* reaching 17%, yet they were outnumbered by *Actinobacteriota* in both moss species – 26% and 24% in *W. fontinaliopsis* and *S. georgicouncinata* respectively. Moreover, *Actinobacteriota* abundance was significantly different between the insects’ and mosses microbiome based on the results of Kruskall-Wallis test (p = 0.009 and p = 0.0023 for S.*georgicouncinata* and *W. fontinaliopsis* respectively). The rest of the microbial community was represented by the phyla with lower abundance fluctuating depending on the sample type – *Firmicutes*, *Cyanobacteria*, *Chloroflexi*, *Acidobacteriota*, *Myxococcota*, *Gemmatimonadota* and the others.

The Unique Insect group exhibited a highly heterogeneous taxonomic composition. However, representatives of *Gammaproteobacteria*, *Alphaproteobacteria*, *Acidobacteriia*, *Actinobacteriia*, *Bacteroidia*, *Clostridia*, and unidentified Bacteria were among the most numerous.

The main difference between the insect and the substrate microbial community at the class level was a significantly higher abundance of *Actinobacteria* in the latter – up to 24% in *W. fontinaliopsis* and 23% in *S. georgicouncinata* (Fig. 4). *Gammaproteobacteria* constituted the major part of *B. antarctica* microbiome collected from *W. fontinaliopsis* (54% under high ornithogenic impact), while the ratio of this group in insects from *Sanionia* was lower (36% in specimens from the locations with low ornithogenic influence) (Fig. 4). The representatives of *Bacilli* were only present in insects’ microbiome samples with the maximum of 24% in *B. antarctica* collected in *S. georgicouncinata*. *Clostridia, Betaproteobacteria* and *Thermoleophilia* were significantly more abundant in substrate samples reaching 5.2%, 6.4% and 9.2% respectively in the *S. georgicouncinata*.

Numerically, most microbial classes either showed a lower abundance in the insect microbiome compared to the corresponding substrate, or their abundance was approximately the same (Figs. S4A, B, C, Figs. S5A, B, C). The classes that prevailed in the insect microbiome collected from the *W. fontinaliopsis* substrate, compared to the *W. fontinaliopsis* microbiome itself, included *Gammaproteobacteria*, *Clostridia*, *Polyangia*, *Babeliae*, *Phycisphaerae*, *Chlamidiae*, and *Fimbriimonadia*. In the microbiome of the insect collected from the *S. georgicouncinata* substrate, the abundance of the class *Bacteroidia* was notably higher compared to the microbiome of the *S. georgicouncinata* substrate itself. Likewise, *Clostridia*, *Gammaproteobacteria*, *Chlamidiae*, *Fimbriimonadia*, and *Phycisphaerae* were more abundandant.

Family level allowed for detection of more fine-scale variation between the unique microbiome of *B. antarctica* and shared with its substrates (Figs. S5A, B, C). Indeed, the members of major families *Yersiniaceae*, *Chitinophagaceae*, *Polyangiaceae*, *Weeksellaceae* and *Fimbriimonadaceae*members and *Peptostreptococcales-Tissierales* group were more abundant in insect microbiota compared to the moss group with up to 108179, 27709, 38789, 34491, 33378 reads, respectively (Fig. S5A). Bacterial families exclusive to the Unique insect microbiota included: *Celulomonadaceae*, *Anaplasmataceae*, *Parvibaculales PS1 clade* and *Akkermansiaceae*. Additionally, substrate-specific differences in the studied microbial community composition were detected, *Azospirillaceae, Chromobacteriaceae, Steroidobacteraceae, Leptolyngbyaceae, Solimonadaceae, Nitrosopumilaceae* and *Puniceispirillales SAR116_clade* members were only found in insects from *S. georgicouncinata* (Fig. S5C).

At the genus level, the taxa that were not identified at the genus level (“*Others*”) comprised the highest ratio in each sample, ranging from 10 % in insects from *W. fontinaliopsis* to 44.7 % in *Sanionia* substrate samples (Fig. 5). The members of the *Yersinia* genus were second most abundant in *B. antarctica* samples – from 0.002 % in *S. georgicouncinata* to 31.9 % in insects from *W. fontinaliopsis*. Additionally, *Chryseobacterium*, *Polaromonas*, *Pseudomonas* and *Luteolibacter* had the highest ratio in microbial communities of insects collected in *W. fontinaliopsis* – averaging at 11.7%, 7.6%, 5.7% and 5.8% respectively (Fig. 5). Meanwhile, *Rhodoferax*, *Carnobacterium*, *Pseudarcicella, Chryseobacterium,* and *Roseiarcus* and peaked in *B. antarctica* collected in *S. georgicouncinata* at 25.9%, 23.8%, 17.9%, 6.1%, 3.5% and on average.

Statistically significant differences were detected in major classes’ distribution between microbial communities under the differential ornithogenic impact. Namely, according to Mann-Whitney test (p-value < 0.05), *Gammaproteobacteria*, *Bacteroidia*, *Verrucomicrobia* and *Bacilli* were significantly enriched in insect samples from *Warnstorfia* collected under high ornithogenic impact, while *Alphaproteobacteria*, *Actinobacteria*, *Cyanobacteria*, *Acidobacteriae*, *Acidimicrobiia*, *Polyangia*, *Planctomyces*, *Thermoleophilia*, and *Gemmatimonadetes* were more abundant in *B. antarctica* from locations with low ornithogenic impact (Fig. 4). However, no significant difference was detected in the *B. antarctica* microbiome from *Sanionia* under the differential ornithogenic impact.

Similarly to the class-level analysis, statistically significant differences (p < 0.05) in microbial taxa distribution were detected between *B. antarctica* samples from *W. fontinaliopsis* under varying ornithogenic impact. Specifically, *Chryseobacterium, Carnobacterium*, *Pseudarcicella* and *Luteolibacter* were more abundant in high-impacted samples, while *Rhodovastum*, *Roseiarcus*, *Mucilaginibacter*, *Granulicella*, *Bradyrhizobium*, *Lacunisphaera*, *Pajaroellobacter*, *Galbitalea*, *Nakamurella*, and *Jatrophihabitans* were enriched in microbiome of *B. antarctica* inhabiting moss under lower ornithogenic influence (Fig. 5, Table S3).

The taxa that contribute most to the variation of microbial community structure were identified and visualized with the PCA (Fig. 6, Table S4). The genera clustered according to their distribution across the different samples’ group: 1. *Yersinia*, *Chryseobacterium*, *Luteolibacter*, *Pseudarcicella*, *Pseudomonas*, *Carnobacterium*, *Rhodoferax* and *Variovorax*; 2. *Roseiarcus*, *Bradyrhizobium*, *Rhodovastrum*, *Mucilaginibacter*, *Pajaroellobacter*, *Granulicella*, *Gabitaleae* and *Lacunisphaera*; 3. *Jatrophihabitans, Dokdonella*, *Nakamurella*, *Gammatimonas*, *Clostridium* and *Mycobacterium*; 4. *Arthrobacter*, *Rhizobacter*, *Oryzihumus*, *Intrasporangium*, *Rhodanobacter*, *Polaromonas* and *Ferrunigibacter* (Fig. 6). This differentiation was supported with the results of Mann-Whitney test (Table S3).

### 3.3. Factors shaping the structure of microbial communities

Following factors had a significant effect on the structure of the studied microbial communities according to the results of Mantel test: ornithogenic impact (r = 0.367, p = 0.0001), sample type (insect or moss, r = 0.074, p = 0.035) and storage solution (r = 0.083, p = 0.011). There was no significant influence of the moss species on the resulting taxonomy of insect or moss microbiome.

Constrained ordination analysis supported this finding, as ornithogenic impact and microbial community origin (insect or substrate) explained the majority of variation – 65% and 31% along the first and the second axis respectively (Fig. 7, Table S5).

### 3.4. The presence of obligate endosymbiotic bacteria in *Belgica antarctica*

Among the most studied obligate endosymbionts of arthropods, only the *Wolbachia* genus was detected in the *B. antarctica* samples. This bacterium has been detected in 2 samples of insects collected in *S. georgicouncinata* and 3 samples of insects from *W. fontinaliopsis*, all ethanol-stored (Table S6). The average abundance of *Wolbachia* genus members in *W. fontinaliopsis* insects microbiome was 0.11%, while this ratio was twice as low in *B. antarctica* collected in *S. georgicouncinata* – 0.06%. All of the samples with the presence of *Wolbachia* were from the locations with high ornithogenic influence, except for the one coming from the low-impacted area with the minimal abundance of this genus.

*Cardinium* and *Rickettsia*, were identified in minor abundance in the studied samples, reaching 4 and 24 reads respectively in insects from *Warnstorfia*. *Arsenophonus* and *Spiroplasma* were not detected in the current study (Table S6).

## 4. Discussion

Microbial communities are recognized to play a crucial role in host adaptation to environmental conditions, which can be of particular importance in hostile Antarctic environment. Insects, particularly members of the family *Chironomidae*, host microorganisms at all stages of their life cycle, yet the information on microbiome of Antarctic terrestrial invertebrates is still scarce. Nowadays, only a few studies have focused on microbiome analysis in representatives of this group: the chironomid *B. antarctica* [Maistrenko et al., 2023], mites *Alaskozetes antarcticus* (Michael, 1903) [Holmes et al., 2019], nematodes *Eudorylaimus antarcticus* (Steiner, 1916) Yeates, 1970 and *Plectus murrayi* Yeates, 1970, unidentified tardigrades [McQueen et al., 2022, 2023], and springtails *Cryptopygus antarcticus* Willem, 1901 and *Friesea antarctica* (Willem, 1901) [Leo et al., 2021]. To our best knowledge to date, microbial communities inhabiting *B. antarctica* have only been analyzed *in silico* with bioinformatic tools and remain insufficiently studied.

Therefore, our study significantly contributes to the current understanding of Antarctic terrestrial insects’ microbial diversity. Both insect and substrate microbiome was analyzed under the differential ornithogenic impact, revealing pronounced differences in between *B. antarctica* and moss microbiome composition, as well as providing insight into the factors shaping the community.

### 4.1. The effect of substrate moss species on *Belgica antarctica* microbiome

We studied the composition of the insect’s microbial communities, as well as the substrate on which the insect larvae lived and fed, namely the mosses *S. georgicouncinata* and *W. fontinaliopsis*. As the diet is known to influence the composition and functioning of the animal microbiome, particularly the insect microbiome [Gohl et al., 2022; Mugo-Kamiri et al., 2024; Adam et al., 2025; Ngando et al., 2025], we aimed at analyzing how decisive the impact of the food source is on the microbiome of *B. antarctica*. The members of Alpha-, Beta-, and Gammaproteobacteria, Actinobacteria, and Bacteroidetes comprised the majority of the microbial diversity, but in distinct proportions depending on the sample type, in both the moss substrates and the insect. The Kruskal-Wallis test revealed a statistically significant difference between the microbial communities of the insects and the moss substrate. The following bacterial classes identified in the current study have been previously reported in the members of *Chironomidae*: *Alphaproteobacteria*, *Actinobacteria*, *Bacteroidia*, *Bacilli*, *Clostridia*, *Gammaproteobacteria*, *Verrucomicrobia*, *Betaproteobacteria*, and *Fusobacteria* [Halpern & Senderovich, 2015]. *Deltaproteobacteria*, *Epsilonproteobacteria*, *Flavobacteria*, *Sphingobacteria*, *Caldilineae*, *Erysipelotrichia*, found by Halpern and Senderovich (2015) were not detected in our samples. The members of *Acidimicrobiia*, *Acidobacteriae*, *Cyanobacteriia*, *Gemmatimonadetes*, *Planctomycetes*, *Polyangia*, and *Thermoleophilia* were identified in *B. antarctica* larvae in our study, yet absent from the previous datasets. This pattern is likely influenced by the available food resources and/or environmental conditions.

To understand which component of the *B. antarctica* microbiome is substrate-derived, we partitioned the OTUs of this insect microbiome into three groups: OTUs common to both the insect and the substrate, OTUs unique to *B. antarctica*, and OTUs common to all insects (Mutual Insect).

The relative abundance of the unique OTUs for *B. antarctica* was negligible (averaging only 1.7%). This portion of the microbiome showed very high heterogeneity among individual samples, suggesting that it does not constitute the core microbiome of *B. antarctica*, but rather represents the unique, individual microbiota of each specimen. This is likely a result of subtle differences during individual larval development and is due to a variation of environment conditions during the long, two-year life cycle of the insect [Sugg et al., 1983; Usher & Edwards, 1984]. Significant portion of the specific *B. antarctica* microbiome diversity was represented by the classes *Acidobacteriia*, *Actinobacteriia*, *Alpha-* and *Gammaproteobacteria*, *Bacteroidia*, and *Clostridia*, as well as unidentified Bacteria.

The OTUs shared by the insect and the substrate quantitatively predominated in the *B. antarctica* microbiome, making up an average of 95% of the entire community. This indicates a fundamental influence of the food source on the *B. antarctica* microbiome. Furthermore, the group of OTUs common to insects inhabiting different substrates (Mutual Insect), which we identified to represent the core microbiome of *B. antarctica*, did not differ taxonomically or statistically from the mutual microbiome of the insects and the mosses. The Mutual Insect group comprised 2930 OTUs with the majority represented by *Gammaproteobacteria, Bacteroidia, Clostridia, Alphaproteobacteria, Polyangia, Acidobacteriae, Actinobacteria, Phycisphaerae*, and *Babeliae* classes. While most OTUs are substrate-derived, the *Belgica* gut performs selective action, altering the ratio of these taxa, and crucially, this portion of the microbiome is not influenced by the moss species.

The question of whether this microbiota is functionally active inside the insect naturally remains open. The detected microbiome, based on the method used (16S rRNA gene sequencing), may include dead and inactive microorganisms from the environment, or transient microorganisms passing through the insect gut. It is most likely that the shared microbiome between *B. antarctica* and the moss substrate consists of both transient and symbiotic microorganisms. Given the limitations of the 16S rRNA amplicon sequencing method, we can only indirectly discuss which members of the community may have a functional role for this Antarctic insect. Specifically, this concerns the taxonomic groups of microorganisms that are enriched in the insect microbiome compared to the substrate microbiome. These classes, which increased in abundance, included *Gammaproteobacteria, Bacteroidia, Clostridia, Polyangia, Babeliae, Phycisphaerae, Chlamidiae*, and *Fimbriimonadia*.

More fine-grained differences between the unique microbiome of *B. antarctica* and shared with its substrate were detected at the family level. Notably, the members of major families *Yersiniaceae, Chitinophagaceae, Polyangiaceae, Weeksellaceae*, and *Fimbriimonadaceaemembers*, and *Peptostreptococcales-Tissierales* group numerically prevailed in insect microbiome compared to the moss.

Members of the *Yersiniaceae* are frequently found in the microbiomes of *Chironomidae* [Rouf & Rigney, 1993; Senderovich & Halpern, 2012, 2013; Halpern & Senderovich, 2015; Sela & Halpern, 2022]. The detoxification role of these bacteria for chironomids, in particular the reduction of the negative impact of heavy metals on the body, is best studied [Laviad-Shitrit et al., 2021; Sela & Halpern, 2022].

The role that species of the *Chitinophagaceae* family may play for chironomids remains unknown, given that these bacteria are characterized by the ability to decompose cellulose and chitin [Rosenberg, 2014], and since this insect can feed on both fungi and dead plant remains [Kozeretska et al., 2022], it is likely that these microorganisms may be involved in *B. antarctica* nutrition.

Members of the *Polyangiaceae* family have been detected in the larvae of several acid-tolerant *Chironomus* species but not in those of acid-sensitive species [Fujii et al., 2023]. It should be noted, although *B. antarctica* larvae can withstand significant pH fluctuations for several weeks [Baust & Lee, 1987; Gantz et al., 2020], this species is often found in acidic environments (pH ≈ 4.0) in nature [Baust & Lee, 1983, 1987]. This may be facilitated by the *Polyangiaceae* associated with the larvae. Bacteria of the *Weeksellaceae* family are also known for other chironomids [Sela & Halpern, 2022]. One possible function that these bacteria may provide for these insects may be resistance to heavy metals. Thus, a resistant to ZnCl_2_ strain of *Chryseobacterium joostei* Hugo et al., 2003 [Senderovich & Halpern, 2013] was isolated from larvae and egg masses of species of the genus *Chironomus*.

The relationship of bacteria of the family *Fimbriimonadaceae* and the group *Peptostreptococcales-Tissierales* with chironomid midges, and insects in general, remains unclear. In addition, there is only isolated evidence (e.g., in several species of saproxylic beetles; Kolasa et al. (2025)) of the association of representatives of *Fimbriimonadaceae* with insects.

Therefore, it is most likely that they contribute to the resistance of the insect to adverse environmental factors and the nutrition of *B. antarctica*. However, there is not enough data to indicate the functions that specific microorganisms may perform within the microbiota of this Antarctic midge.

### 4.2. The effect of ornithogenic factor on *Belgica antarctica* microbiome

The microbiome of *B. antarctica* was also analyzed in the context of ornithogenic impact, as the expansion of the *P. papua* population due to climate change is an important factor affecting Antarctic terrestrial ecosystems [Parnikoza et al., 2018, Prekrasna-Kviatkovska et al., 2024]. The moss substrate from which the *B. antarctica* larvae were collected was differentially affected by birds. We divided the intensity of this impact into three levels (low, medium, and high); this scale corresponded to the presence of only skua nests (low), the establishment of *P. papua* colonies since 2019 (medium), and the long-term *P. papua* nesting activity commencing in 2007 (high). Kruskal-Wallis testing and NMDS analysis indicated that the composition of the *B. antarctica* microbial communities is significantly influenced by the intensity of the ornithogenic factor.

The ornithogenic impact on substrates is primarily associated with environmental eutrophication, which results from the introduction of a significant amount of nitrogen, phosphorus, and carbon compounds. Consequently, the metabolic transformation of nitrogen compounds (e.g., ammonification) can lead to an increase in the substrate’s pH. The previous chemical analysis of the moss substrate on Galindez Island revealed that the concentrations of PO₄³⁻ and dissolved inorganic phosphorus (DIP), as well as NH₄⁺, NO₂⁻, NO₃⁻, and the total content of dissolved inorganic nitrogen (DIN), were several tens of times higher in mosses exposed to high ornithogenic influence, and pH of the substrate rose from acidic to neutral [Prekrasna-Kviatkovska et al., 2024]. In addition to enriching the environment with nitrogen- and phosphorus-containing compounds, seabirds contribute to the transfer of heavy metals, including Cu, Zn, Pb, and Hg from the ocean to Antarctic terrestrial ecosystems [Castro et al., 2021; Soares et al., 2024]. Moreover, birds significantly contaminate the substrate with their gut microbiota [Prekrasna-Kviatkovska et al., 2024], which may decline under contrasting environmental conditions or partially adapt [Grzesiak et al., 2020]. On the other hand, the ornithogenic impact on the substrate microbiome manifests in an alteration of environmental conditions, which creates a selective advantage for the development of different groups of microorganisms than those that previously dominated.

The rising intensity of the ornithogenic impact, observed at certain sites of the current study, contributed to the shift in microbial communities composition. Indeed, *Gammaproteobacteria, Bacteroidia, Verrucomicrobia*, and *Bacilli* were significantly enriched in insect samples collected from highly impacted samples. Among genera, *Chryseobacterium, Carnobacterium*, *Pseudarcicella*, and *Luteolibacter* prevailed in the samples from samples with high ornithogenic impact. The genus *Chryseobacterium* is frequently associated with the intestines of animals, including penguins. For instance, a proteolytic *Chryseobacterium* sp. was identified in the guano of *Pygoscelis adeliae* (Hombron & Jacquinot, 1841) [Grzesiak et al., 2020] and in the penguin-disturbed areas of the tall moss turf subformation [Prekrasna-Kviatkovska et al., 2024]. *Chryseobacterium* was shown to harbour a variety of nitrogen metabolism and transport genes [Jung et al., 2023], as well as proteases (e.g., keratinase) that can enhance degradation of protein-rich substances from penguin guano [Bokveld et al., 2021]. *Carnobacteriaceae* were previously found in high abundance dominant in actively penguin-colonized soils [Kim et al., 2012]. Similarly to *Chryseobacterium*, *Carnobacterium* had high activity in protein degradation and toxin tolerance. *Carnobacterium* was recently reported from the insect gut for the first time, signaling a potentially significant role in metabolism and stress resistance of a host [Braglia et al., 2025]. The members of *Luteolibacter* genus have been shown to possess vast mechanisms for carbohydrate degradation and urea transformation under elevated pH levels [Pascual et al., 2017], which can be beneficial for the host under ornithogenic impact. Additionally, heavy metal tolerance was reported for *Chryseobacterium* from *Anopheles arabiensis* gut [Singh et al., 2024], and for *Carnobacterium* [George et al., 2021, Prieto-Fernández et al., 2024].

The shift in the insect’s microbiome due to the ornithogenic factor occurs as a consequence of the modification of the trophic base. Since the substrate is chemically and microbiologically modified, this inevitably reflects on the insect’s microbiome. However, certain taxa were quantitatively dominant in the *B. antarctica* microbiome from the highly affected sites compared to both the microbiomes of the highly affected sites and the microbiome of *B. antarctica* from the sites with low ornithogenic impact. This might suggest that the larval gut mediates the selective pressure for the taxa that contribute to host adaptation. Therefore, significantly higher abundance of *Chryseobacterium*, *Carnobacterium*, and *Luteolibacter* in larvae from highly-impacted sites might be a result of their adaptation to the conditions, induced by ornithogenic factor.

### 4.3. Comparison of our results with previous data

At both the class and genus levels, our data revealed markedly higher taxonomic diversity compared with Maistrenko et al. (2023): 14 classes versus seven, and 30 genera versus 13. None of the five genera that Maistrenko et al. (2023) classified as potential contaminants in their samples were detected in our dataset. The observed differences between the two studies might have arisen from the different geographic location of the sampling sites (Palmer Archipelago in Maistrenko et al. (2023) and Argentine islands Archipelago in our study), which potentially implies varying substrate type, ornithogenic impact and the other environmental factors’ influence. Moreover, the study design differed substantially, both in terms of the insect developmental stages used in the analysis (larvae in the current study versus imagos and larvae in Maisternko et al. (2023)). and in terms of metagenomic analysis methodology. Indeed, the earlier datasets used by Maistrenko et al. (2023) were produced for the purposes other than microbiome analysis, which may have contributed both to contamination with microorganisms not characteristic of *B. antarctica* or its habitat, and to the loss of bacterial sequences, particularly in cases of low bacterial abundance.

Only three genera: *Yersinia*, *Pseudomonas*, and *Arthrobacter* were mutual between the two studies, and are likely components of the core microbiome of *B. antarctica*. Interestingly, *Pseudomonas* was also found in the Antarctic-invasive and closely related *Chironomidae E. murphyi* [Brayley et al., 2025a]. Members of the genera *Yersinia* and *Pseudomonas* are known to potentially contribute to heavy metal resistance. Namely, *Yersinia nurmii* Murros-Kontiainen et al., 2011 (resistant to Pb) and *Pseudomonas geniculata* (Wright, 1895) Chester, 1901 (resistant to Zn) were among numerous bacterial taxa isolated from the egg masses of *Chironomus transvaalensis* Kieffer, 1923 [Sela & Halpern, 2022]. Sela et al. (2021) suggested that in *Chironomus ramosus* Chaudhuri, Das & Sublette, 1992 *Pseudomonas* sp. contributes to metabolic pathways responsible for pollutants’ degradation, for example As. It is also well established that *Pseudomonas* exhibits detoxification activity, including phenol degradation, aromatic hydrocarbon oxidation, and participates in the bioremediation of contaminated environments [Williams & Sayers, 1994; Jõesaar et al., 2017; Sela & Halpern, 2022]. The *Yersinia* abundance in chironomid larvae can increase in response to elevated levels of Cu and Cr in the environment [Laviad-Shitrit et al., 2021], potentially contributing to the host’s adaptation to these metals. The interaction between chironomids and *Arthrobacter* is poorly studied, but it is likely also involved in detoxification, as members of this genus are capable of degrading the chlorotriazine herbicide atrazine [Cai et al., 2003] and the previously widely used plasticizer, butyl benzyl phthalate ester [dos Santos Morais, 2020].

Unlike many other *Chironomidae*, *B. antarctica* larvae inhabit various plant substrates or soil rather than aquatic environments. Furthermore, their small body size and ability to occupy various substrates likely contribute to the instability of their microbiome (Maistrenko et al. (2023). Thus, it can be assumed that the microbiome of chironomids, even at the class level, is strongly influenced by environmental conditions, geographic region, or ecological niche. This is confirmed by the differences we found at this taxonomic level even among individuals of the same species, collected both from different moss substrates and from sites with varying levels of ornithogenic impact. And this is consistent with the hypothesis proposed by Laviad-Shitrit et al. (2021), according to which, a chironomid larvae maintain a consortium of diverse bacterial species, some of which may proliferate in response to environmental conditions and thereby contribute to the host’s survival under changing environmental conditions.

The differences between our study and the previous data on Antarctic *Chironomidae* microbial diversity highlight the necessity of additional sampling effort, which would allow for comparison between the different regions uncovering potential local adaptations of terrestrial invertebrates.

### 4.4. The first report of *Wolbachia* infection in *Belgica antarctica* populations

It is also important to highlight the detection of *Wolbachia* infection in some of the tested samples. *Wolbachia pipientis* Hertig, 1936 is one of the most widespread symbiotic bacteria present across invertebrates: this bacterium infects approximately half of all terrestrial arthropod species [Weinert et al., 2015; Charlesworth et al., 2019] outside Antarctica [Serga et al., 2024]. *Wolbachia* is a symbiont with a wide range of phenotypic effects on different host species, spanning mutualistic, symbiotic, and parasitic relationships [Kaur et al., 2021; Serga et al., 2021; Mioduchowska et al., 2023]. This bacterium is typically transmitted through the maternal germline along vertical transmission [Werren et al., 2008]. The alternative transmission ways, such as hybrid introgression and codivergence, horizontal transfer, are also reported by Baldo et al. (2008) and Raychoudhury et al. (2009). Although the phenomenon of horizontal transfer with totally predominant vertical transmission is known for *Wolbachia*, and events of introduction of new species are quite frequent in Antarctica [Vega et al., 2020; Hughes et al., 2023], obviously, this way of *Wolbachia* spreading cannot be considered effective.

There are several studies that have shown the absence of *Wolbachia* in *B. antarctica* [Holmes et al., 2019; Maistrenko et al., 2023; Serga et al., 2024]. This endosymbiont has also not been found in populations from Signy Island (South Orkney Islands, Antarctica) of *E. murphyi* [Brayley, 2025b], a close-related to the genus *Belgica* [Allegrucci et al., 2012]. One of the hypotheses proposed by Serga et al. (2024) suggests that this may be explained by the tectonic isolation of the Antarctic. However, this is unlikely to represent the primary cause, as *Wolbachia* Supergroups A and B, which infect arthropods, are reciprocally monophyletic and and diverged from their last mutual ancestor around 217 million years ago [Liu et al., 2023], whereas Antarctica became fully separated from South America and Australia only approximately 30 million years ago [van den Ende et al., 2017]. Also, Serga et al. (2024) have proposed several other hypotheses that may explain why no clear evidence of *Wolbachia* has been found in Antarctic terrestrial invertebrates: phylogeography and demographic events in the host populations negatively influence *Wolbachia* spread and maintenance in Antarctica; сold temperatures or other environmental extremes have negative direct effects on *Wolbachia* survival or influence parameters controlling its spread in populations of arthropods in Antarctica; *Wolbachia* infection is too costly for Antarctic invertebrates; and one of the most compelling, especially in the context of this work, is insufficient research.

The bioinformatics search provided by Serga et al. (2024) shows doubtful weak evidence (only one sample with total coverage <0.01% of its genome of *Wolbachia*), suggesting that *Wolbachia* might be present in one sample of *B. antarctica*. It is important to note that direct PCR testing with applying only two pairs of primers (to genes *16S rRNA* and *wsp*) did not confirm *Wolbachia* infection in this species [Maistrenko et al., 2023]. It is necessary to note that the study by Rodrigues et al. (2023) aimed to detect this endosymbiont in 58 Collembola species worldwide, employing six primer pairs that targeted the genes *coxA*, *dnaA*, *fbpA*, *ftsZ*, *gatB*, and *16S rDNA*. Not all primer sets successfully amplified every sample, and the detection of a single genetic marker among those tested was considered sufficient to confirm infection. Due to the possible long Antarctica isolation, as well as the cold conditions in this region, it is possible that if *Wolbachia* is present in invertebrate populations in this area, it may have changed significantly. Therefore, for its detection by PCR, more primers to different bacterial genes should be used.

According to our results, *Wolbachia* was still present in 5 out of 33 studied samples of this wingless midge from Galindez Island, which likely represents the first confirmed detection of this endosymbiont in any of the Antarctic terrestrial invertebrates. This raises the question of the possible role of *Wolbachia* this endosymbiont in *B. antarctica*, and also indicates the necessity for more detailed studies of endosymbiotic bacteria of arthropods of the sixth continent.

### 4.5. On the potential infection of other obligate endosymbionts in *Belgica antarctica*

Unlike *Wolbachia*, the other phenotype-modifying endosymbionts, *Cardinium* and *Rickettsia*, were detected only in minor abundance in the studied *B. antarctica* samples, while *Arsenophonus* and *Spiroplasma* (also not found by Maistrenko et al., 2023) were not found.

*Rickettsia*, a member of the same family as *Wolbachia* (Rickettsiaceae), infects approximately 24% of terrestrial arthropods [Weinert et al., 2015]. It is known that *Rickettsia* in some cases (e.g., *Bemisia tabaci* (Gennadius, 1889)) can contribute to the host species’ survival during changes in temperature, although to an increase [Brumin et al., 2011]. On the other hand, the example of *Rhipicephalus sanguineus* Latreille, 1806 ticks shows that infection with this bacterium increases mortality both at low and high temperatures [Socolovschi et al., 2012]. Therefore, its effects on the host’s thermal tolerance appear to be species-specific. Interestingly, *Rickettsia* was recorded in Antarctica in a species closely related to the *B. antarctica*, namely *E. murphyi*, which is invasive to this region [Brayley et al., 2025a, b]. This bacterium exhibited one of the highest abundances among the entire microbiome [Brayley et al., 2025a]. The results obtained are noteworthy because, unlike *B. antarctica*, *E. murphyi* reproduces by parthenogenesis [Cranston, 1985; Bartlett et al., 2019], and *Rickettsia* can induce it in some hosts [Ma & Schwander, 2017].

*Cardinium* is an endosymbiotic bacterium that infects more than 13% [Weinert et al., 2015] of arthropod species, transmitted mainly transovarially [Weinert et al., 2015; Hubert et al., 2021]. *Cardinium* affects the expression of host genes, including pheromone genes, thus providing itself with an advantage in inheritance [Hubert et al., 2021]. It can cause cytoplasmic incompatibility and feminization [Chigira & Miura, 2005; Doremus et al., 2022]. It has been demonstrated that the phenotypic effects of *Cardinium* are temperature-dependent [Doremus et al., 2019] and are associated with a longer developmental time, the longest of which were identified in arthropods from polar regions [Haghshenas-Gorgabi et al., 2023].

According to various estimates, different *Spiroplasma* members infect from 7 to over 11% of terrestrial arthropod species [Duron et al., 2008; Kakizawa et al., 2022]. This endosymbiont can enhance the insects’ resistance to parasitoid wasps, parasitic nematodes, and fungal pathogens [Lo et al., 2013]. Previous studies also indicate that *Spiroplasma* affects its hosts under various stressors, including low temperatures [Konai et al., 1996; Gasparich, 2010].

Representatives of the genus *Arsenophonus* are estimated to infect less than 5% of terrestrial arthropod species [Duron et al., 2008]. This endosymbiont is known to cause a mele-killing phenotype [Werren et al., 1986; Ferree et al., 2008; Garrido-Bautista et al., 2024]. However, the effects of temperature on symbiont-host interactions for *Arsenophonus* remain poorly understood.

It should be emphasized that the absence or low abundance of these endosymbionts in the studied samples could putatively arise from the bias of 515F-806R barcode universal primers used in the current study for microbial community profiling. It is accepted that any primer pair targeting a specific variable region of 16S rRNA might exhibit bias towards certain microbial taxa while excluding others [Wasimuddin et al., 2020]. Even though this primer pair can detect endosymbiotic bacteria [Nowak et al., 2025], the failure to detect *Arsenophonus* and *Spiroplasma* endosymbionts, does not allow speculating about their absence in *B. antarctica* larvae in the current study. Further research are also required to confirm the infection of *B. antarctica* with the endosymbionts *Cardinium* and *Rickettsia*.

## 5. Conclusions

The structure of microbial communities is shaped by the ornithogenic influence level. The significant differences observed between the *B. antarctica* bacterial communities and their associated *W. fontinaliopsis* substrate under high ornithogenic impact suggest that the larval microbiome is not a simple reflection of the surrounding substrate but rather a consequence of the selection of microorganisms that contribute to the host adaptation.

We identified the groups of microorganisms that constitute the so-called core microbiome of *B. antarctica*, which remains stable and consistently present in the insect regardless of the moss substrate on which it occurs. However, the substrate microbiome is the primary source of the microbial diversity associated with *B. antarctica*.

For the first time, the presence of the endosymbiont *Wolbachia* in *B. antarctica* has been detected, with the bacterium found in 5 out of 33 tested samples. This is the first confirmation of this endosymbiotic bacterium among Antarctic invertebrates.

## Abbreviations

CAP: constrained ordination
NMDS: non-metric multidimensional scaling
OTU: Operational taxonomic unit
PCA: principal component analysis
PD: phylogenetic distance

## Supplementary information

This manuscript contains 6 Supplementary Tables (Tables S1–S6) and 5 Supplementary Figurers (Figs. S1–S5).

## Acknowledgements

Not applicable.

## Authors’ contributions

PK performed investigation, visualisation, data curation, and Writing–original draft. MP performed methodology, investigation, statistical analysis, visualisation, data curation, and Writing–original draft. YeP-K performed methodology, investigation, data curation, validation, and Writing–original draft. AP performed samples collection and Writing–review and editing. IK performed conceptualization, validation, project administration, supervision, Writing–original draft.

## Funding

This study was carried out in the framework of the Ukrainian State Special-Purpose Research Program in Antarctica for 2011–2025. Pavlo Kovalenko is supported by the scientific research project of the National Academy of Sciences of Ukraine “Ecological indication of biotopes conditions using model biota representatives under anthropogenic transformation of ecosystems” (Grant No. 0125U000528).

## Data availability

The sequences obtained in this study have been deposited to the NCBI database under the accession number PRJNA1374202.

## Declarations

### Ethics approval and consent to participate

Not applicable.

### Consent for publication

Not applicable.

### Competing interests

The authors state no competing interests.

## Supplementary information

**Table S1.**
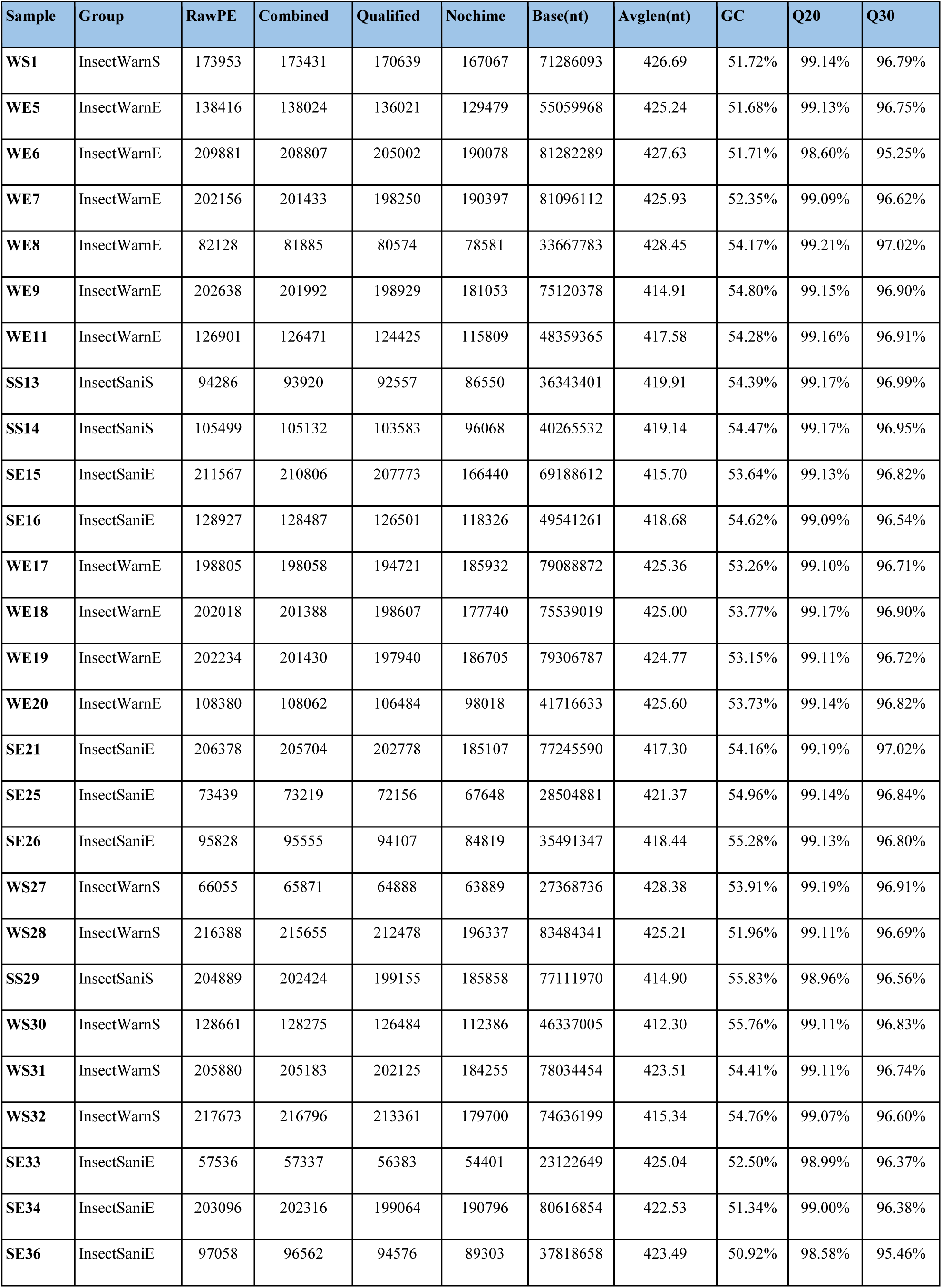
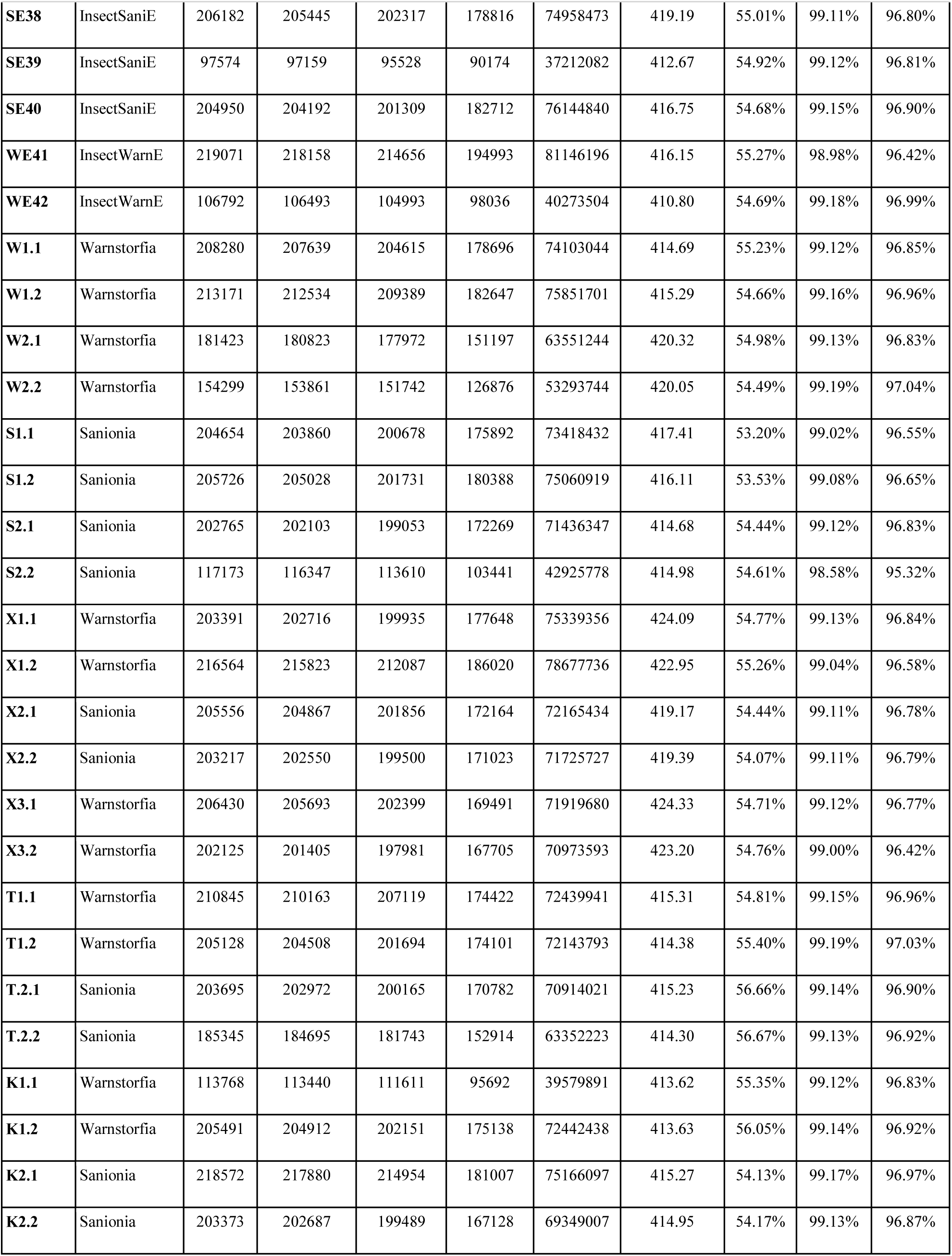
Sequencing statistics.

**Table S2.**
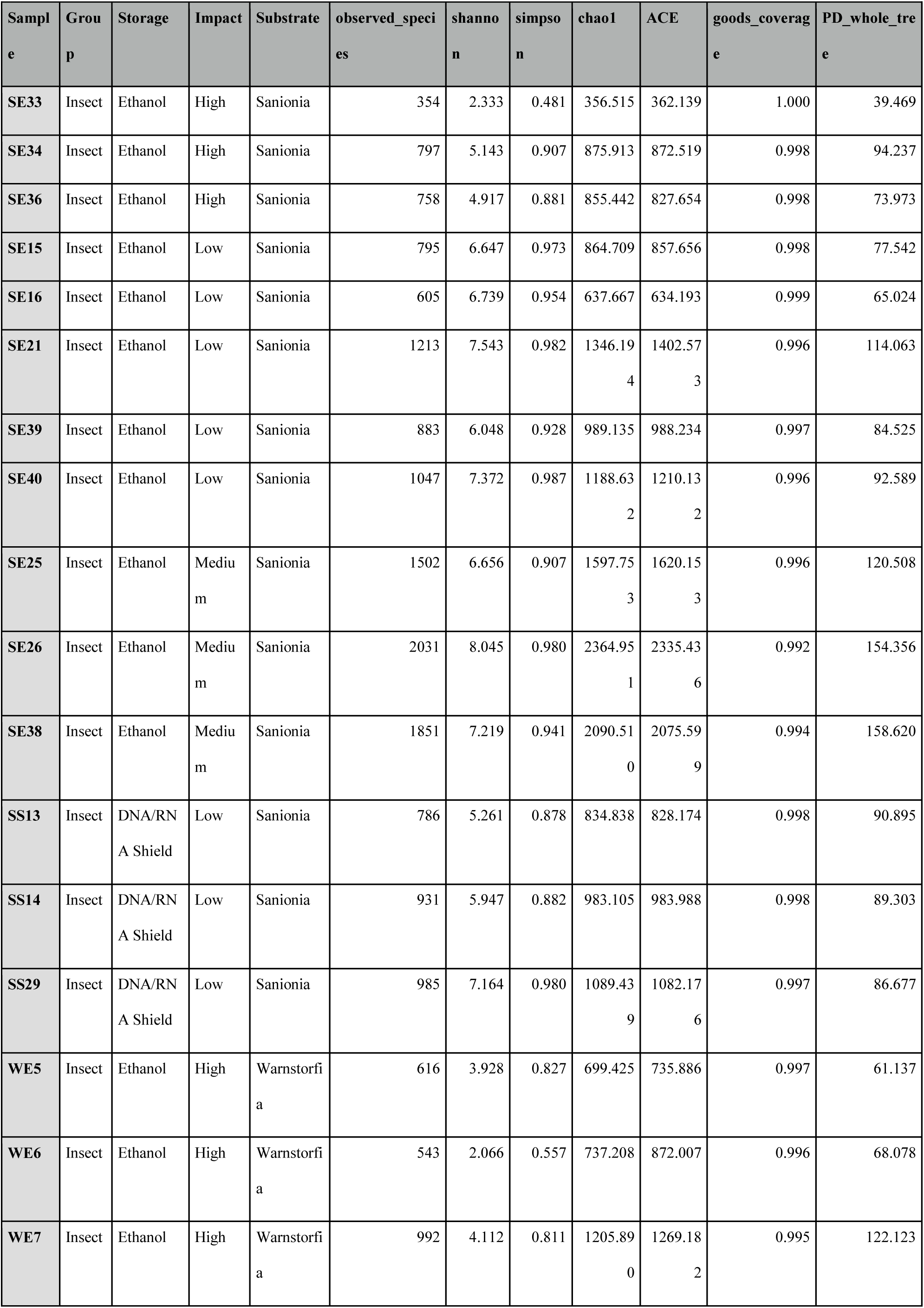
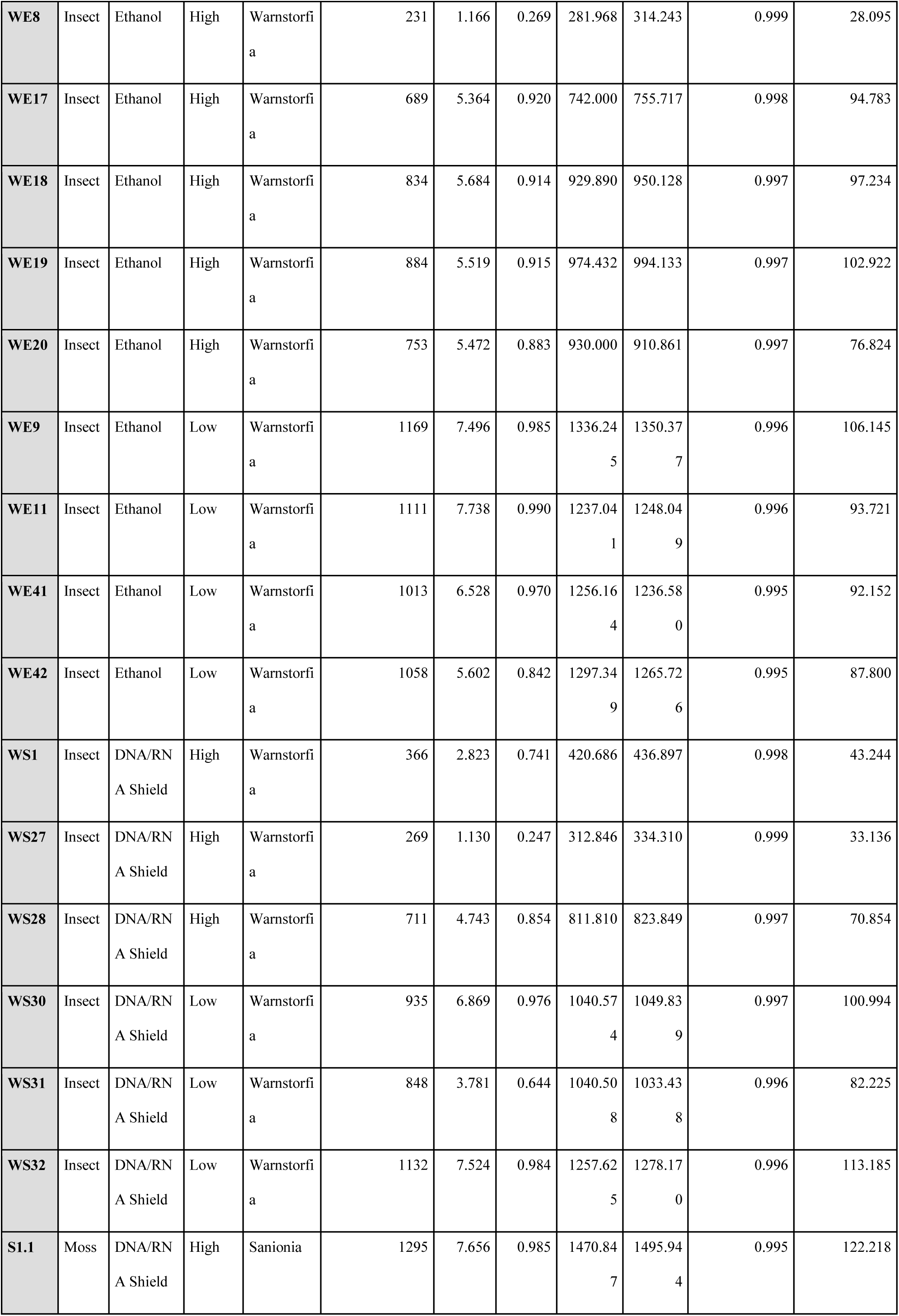
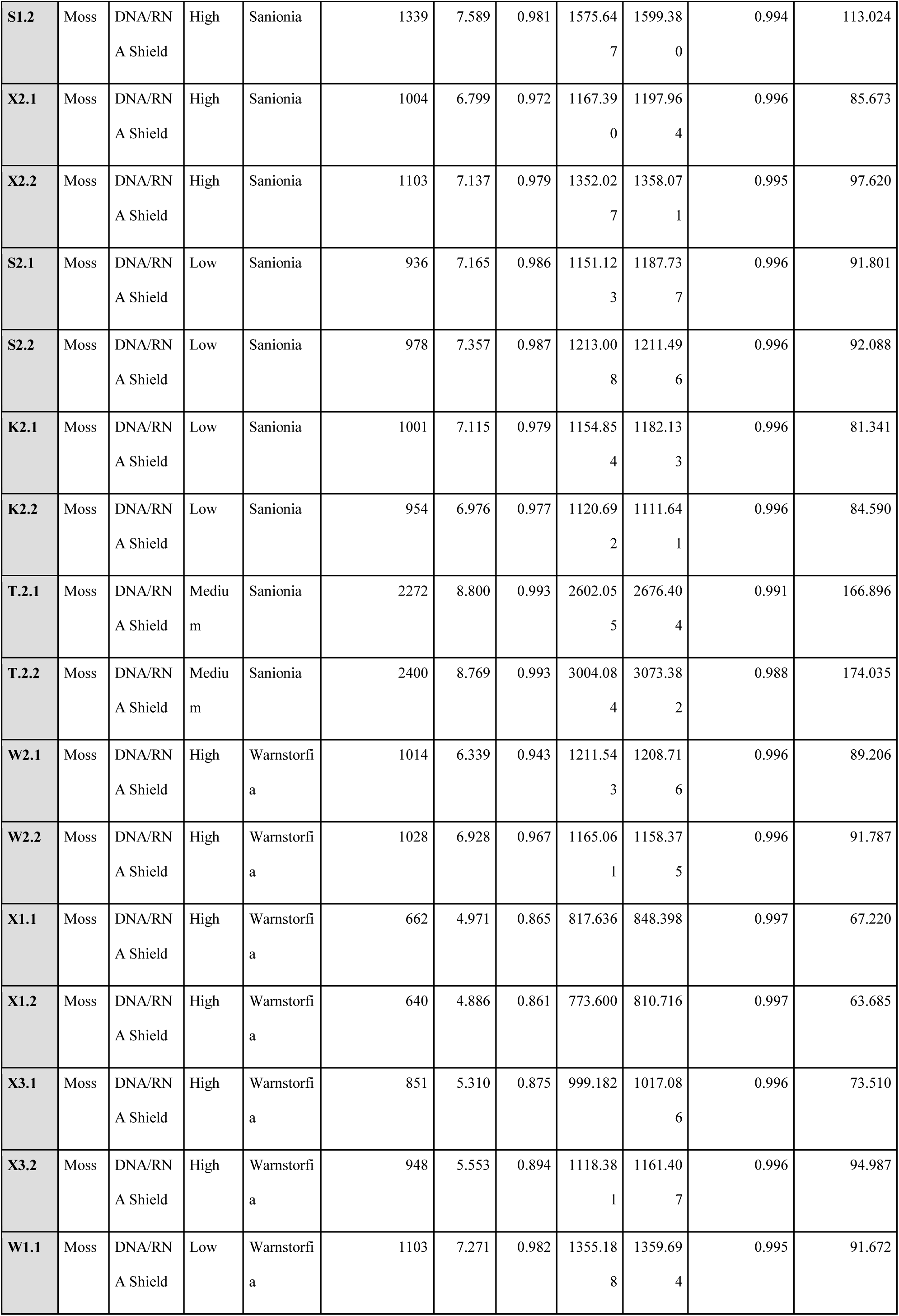
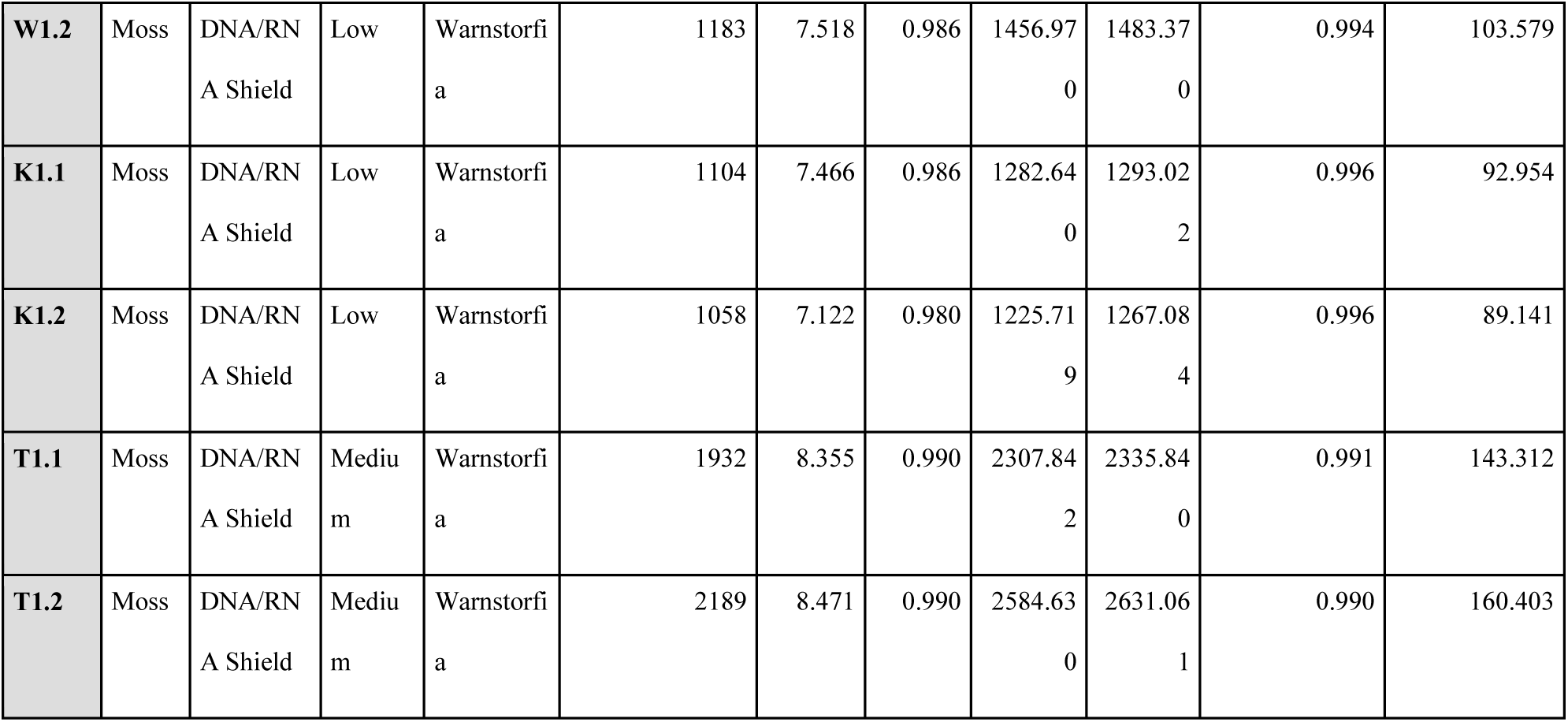
Microbial communities diversity metrics.

**Table S3.**
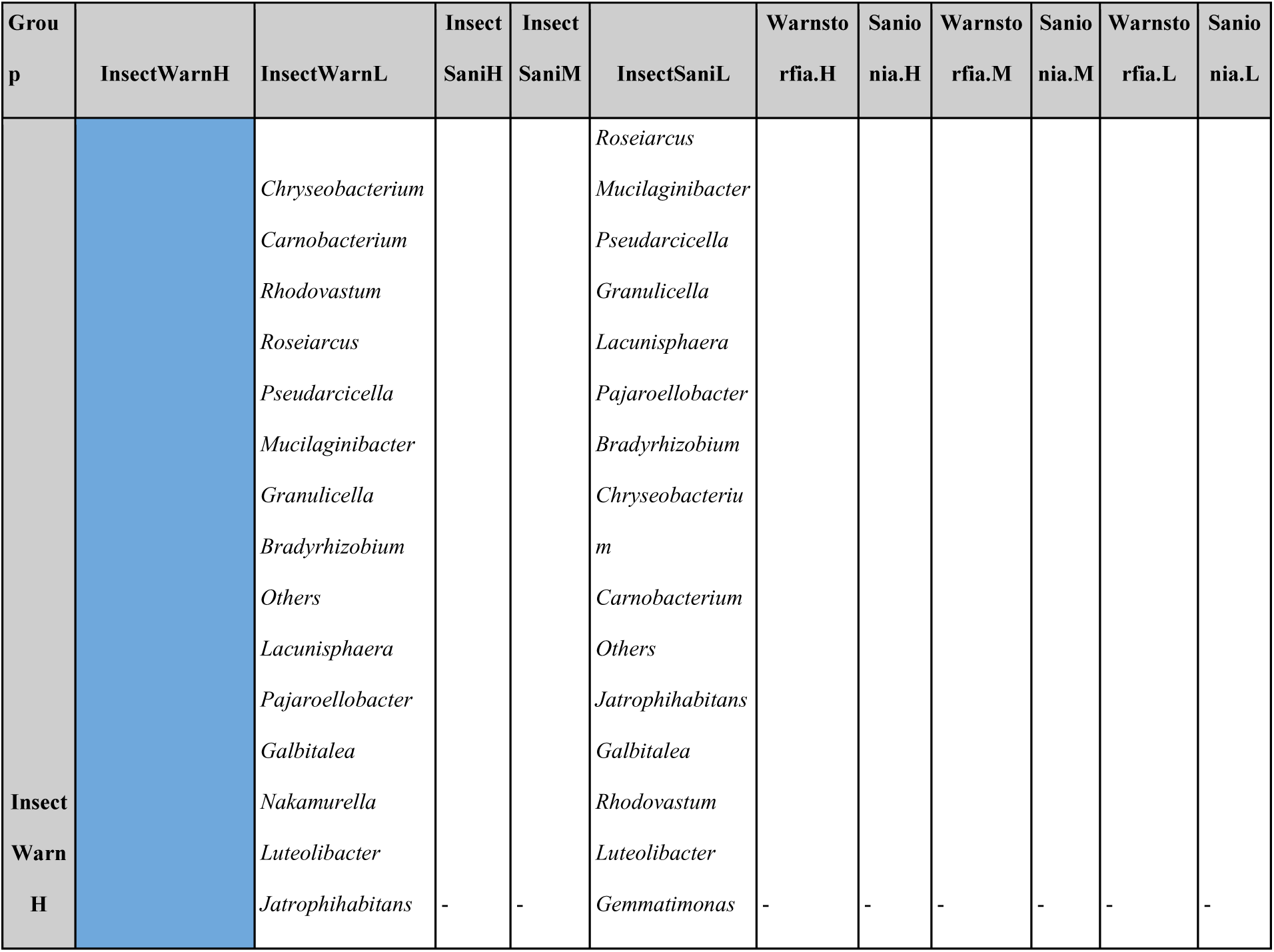
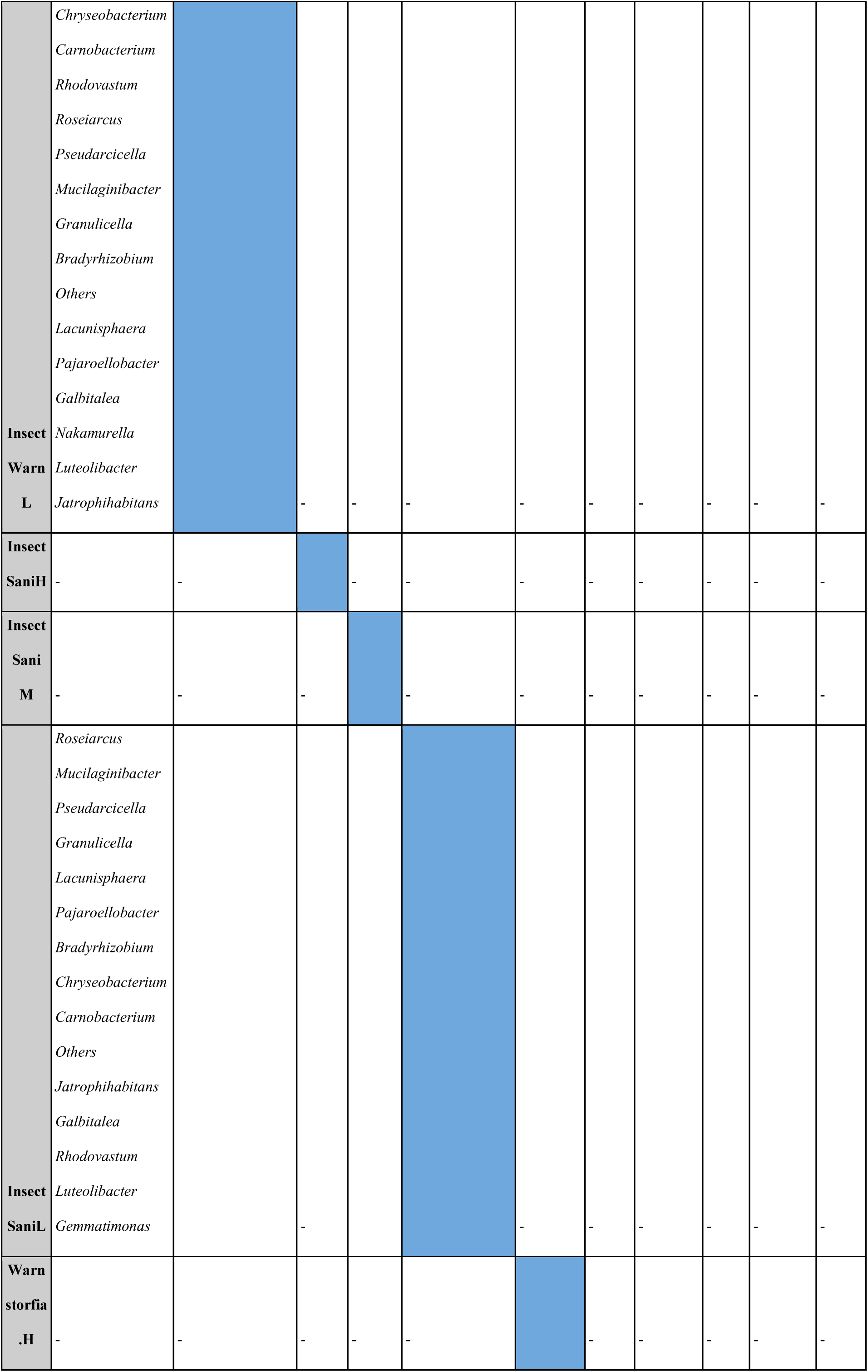
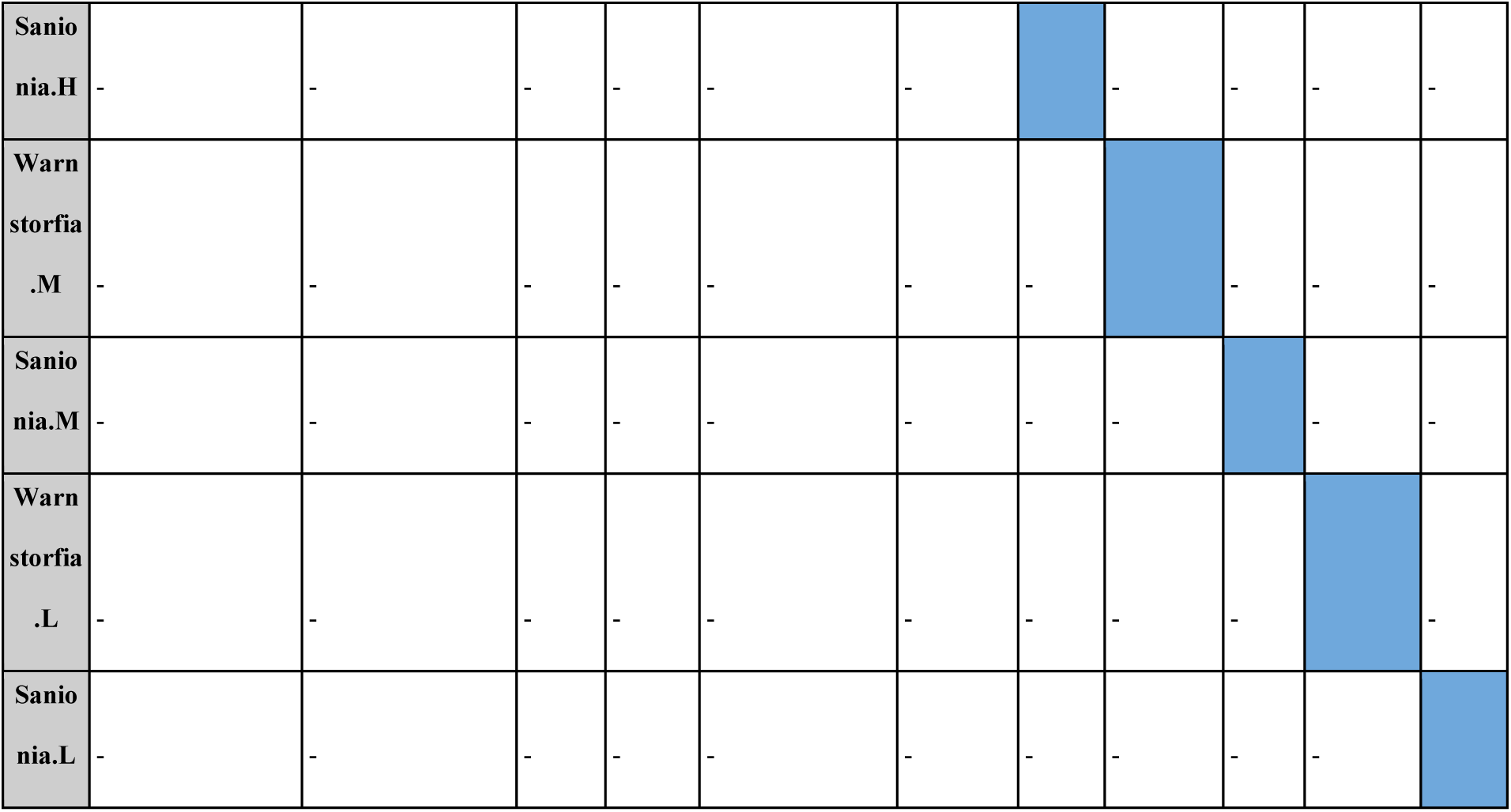
The genera with significantly different distribution across the samples’ types (Mann-Whitney test, p<0.05)

**Table S4.** Eigenvalues of the principal components (PC) that explain the most variance across microbial composition of different samples.

| PC summary | PC1 | PC2 |
| --- | --- | --- |
| **Eigenvalue** | 6,915 | 5,012 |
| **Proportion of variance** | 23,05% | 16,71% |
| **Cumulative proportion of variance** | 23,05% | 39,76% |

**Table S5.** CAP eigenvalues (accumulated, constrained) and their contribution to the squared Bray distance.

|  | CAP1 | CAP2 |
| --- | --- | --- |
| **Eigenvalue** | 2.5630 | 1.2800 |
| **Proportion Explained** | 0.6669 | 0.3331 |
| **Cumulative Proportion** | 0.6669 | 1.0000 |

**Figure S1.**
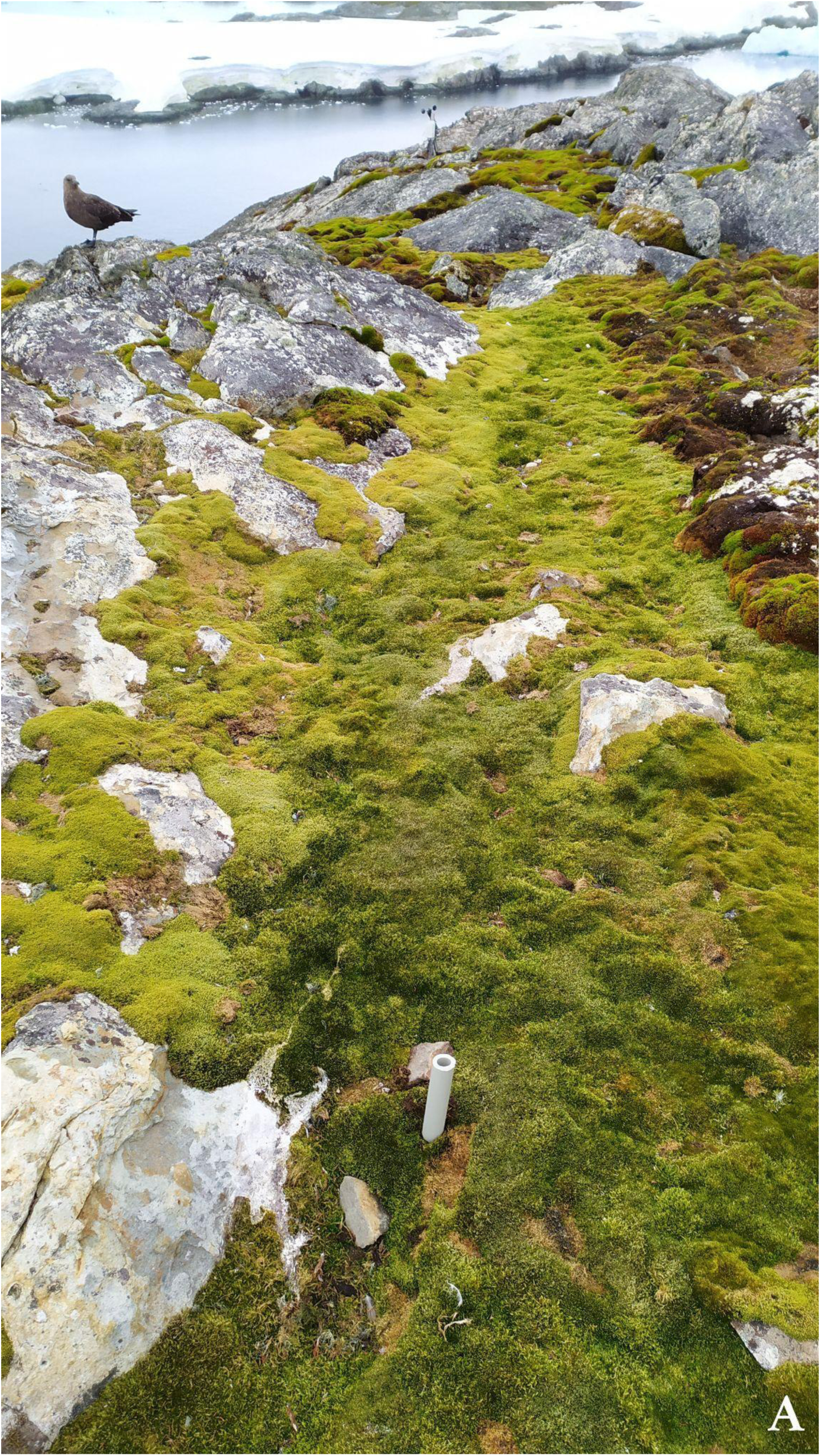
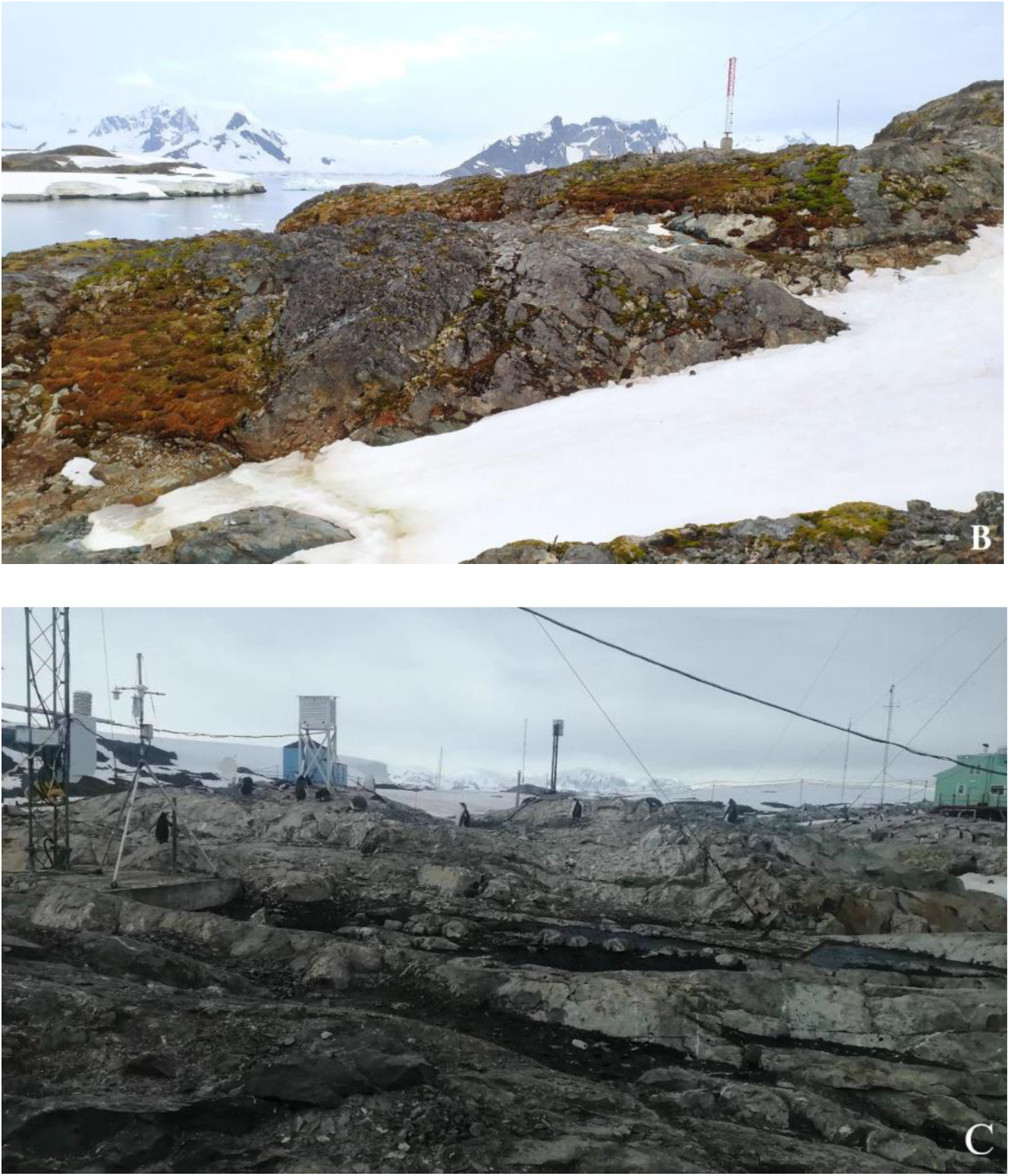
Photos of locations with varying levels of ornithogenic impact where samples were collected: A – low, B – medium, C – high

**Figure S2.**
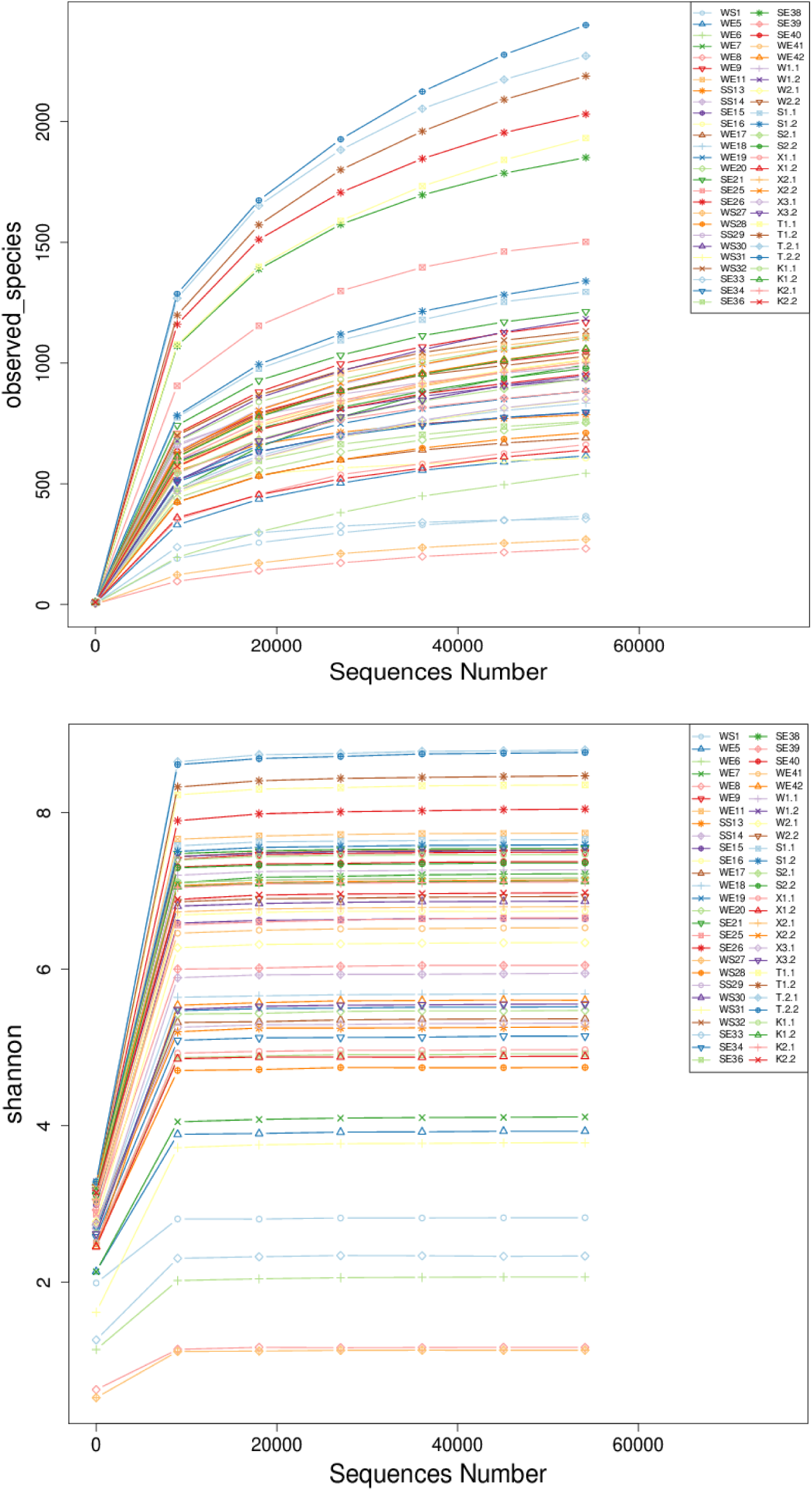
Clustering of different samples’ groups based on the NMDS analysis (squares – medium, crosses – high and circles – low ornithogenic impact). InsectSaniS – individuals of *Belgica antarctica* from the *Sanionia georgicouncinata* substrate, fixed in a DNA/RNA Shield solution; InsectSaniE – specimens of *Belgica antarctica* from the *Sanionia georgicouncinata* substrate, fixed in 96% ethanol; InsectWarnS – individuals of *Belgica antarctica* from the *Warnstorfia fontinaliopsis* substrate, fixed in a DNA/RNA Shield solution; InsectWarnE – specimens of *Belgica antarctica* from the *Warnstorfia fontinaliopsis* substrate, fixed in 96% ethanol; Sanionia – *Sanionia georgicouncinata* substrate; Warnstorfia – *Warnstorfia fontinaliopsis* substrate

**Figure S3.**
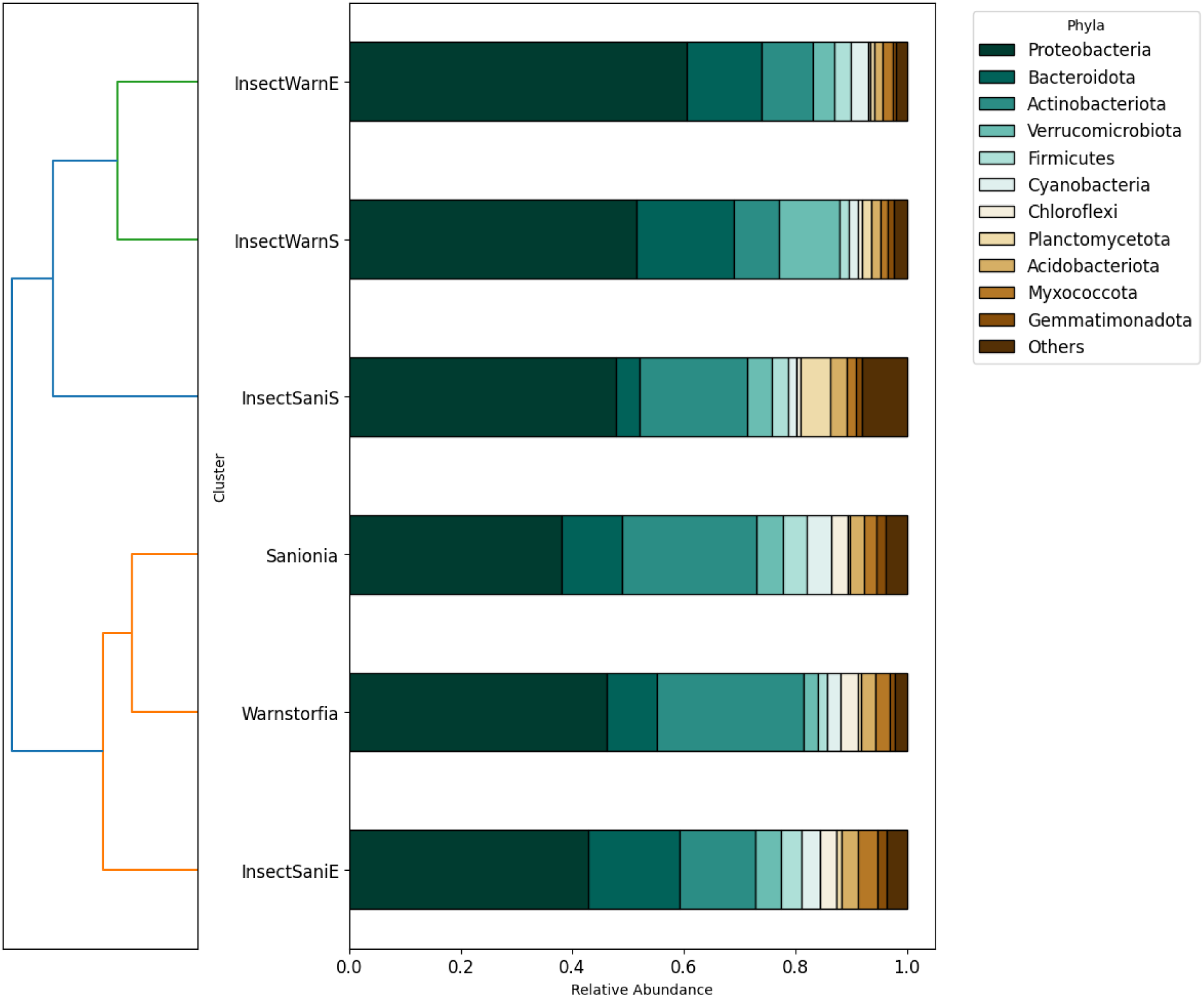
Clustering of different samples’ groups based on the Weighted Unifrac Distance. InsectSaniS – *Belgica antarctica* specimens from the *Sanionia georgicouncinata* substrate, fixed in a DNA/RNA Shield solution; InsectSaniE – *Belgica antarctica* individuals from the *Sanionia georgicouncinata* substrate, fixed in 96% ethanol; InsectWarnS – *Belgica antarctica* individuals from the *Warnstorfia fontinaliopsis* substrate, fixed in a DNA/RNA Shield solution; InsectWarnE – larvae of *Belgica antarctica* from the *Warnstorfia fontinaliopsis* substrate, fixed in 96% ethanol; Sanionia – *Sanionia georgicouncinata* substrate; Warnstorfia – *Warnstorfia fontinaliopsis* substrate

**Figure S4.**
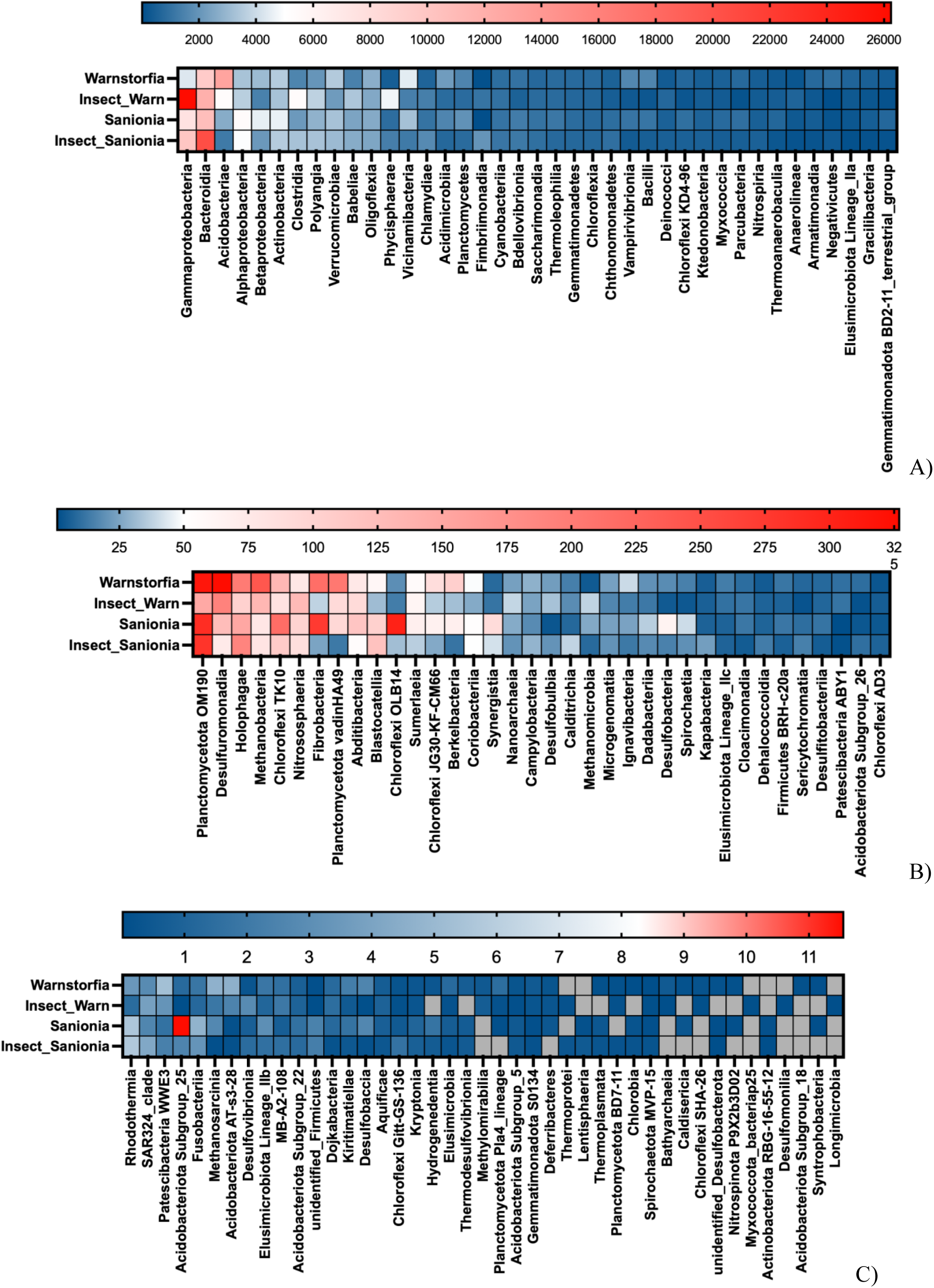
Taxonomic distribution (number of reads) of microbial communities at the class level in different types of samples (Warnstorfia – *Warnstorfia fontinaliopsis* moss, Sanionia *- Sanionia georgicouncinata moss,* InsectWarn – *Belgica antarctica* from the *Warnstorfia fontinaliopsis,* InsectSanionia – *Belgica antarctica* from the *Sanionia georgicouncinata*): A) average abundance < 26230 reads, B) average abundance < 326 reads , C) average abundance < 11.5 reads

**Figure S5.**
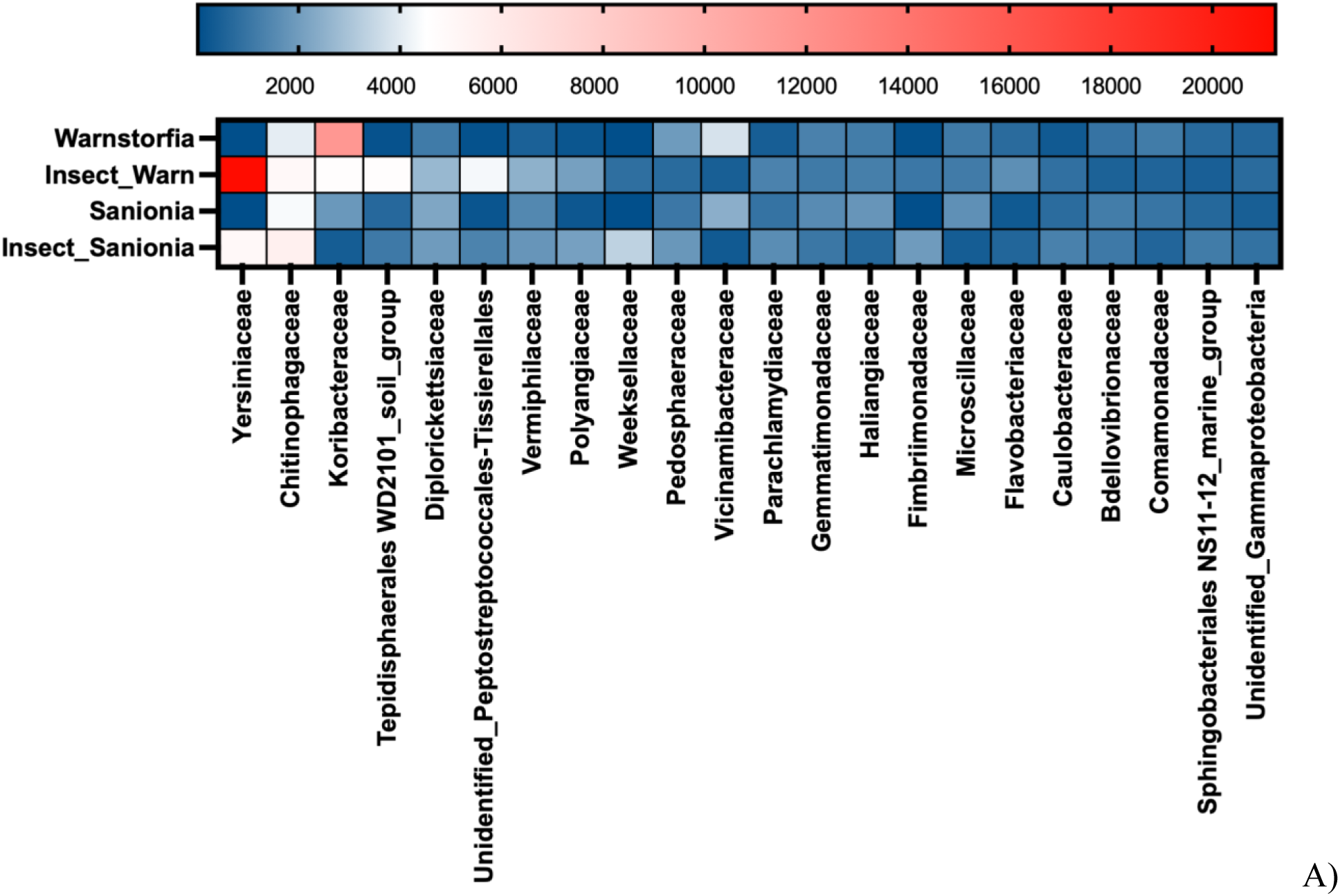
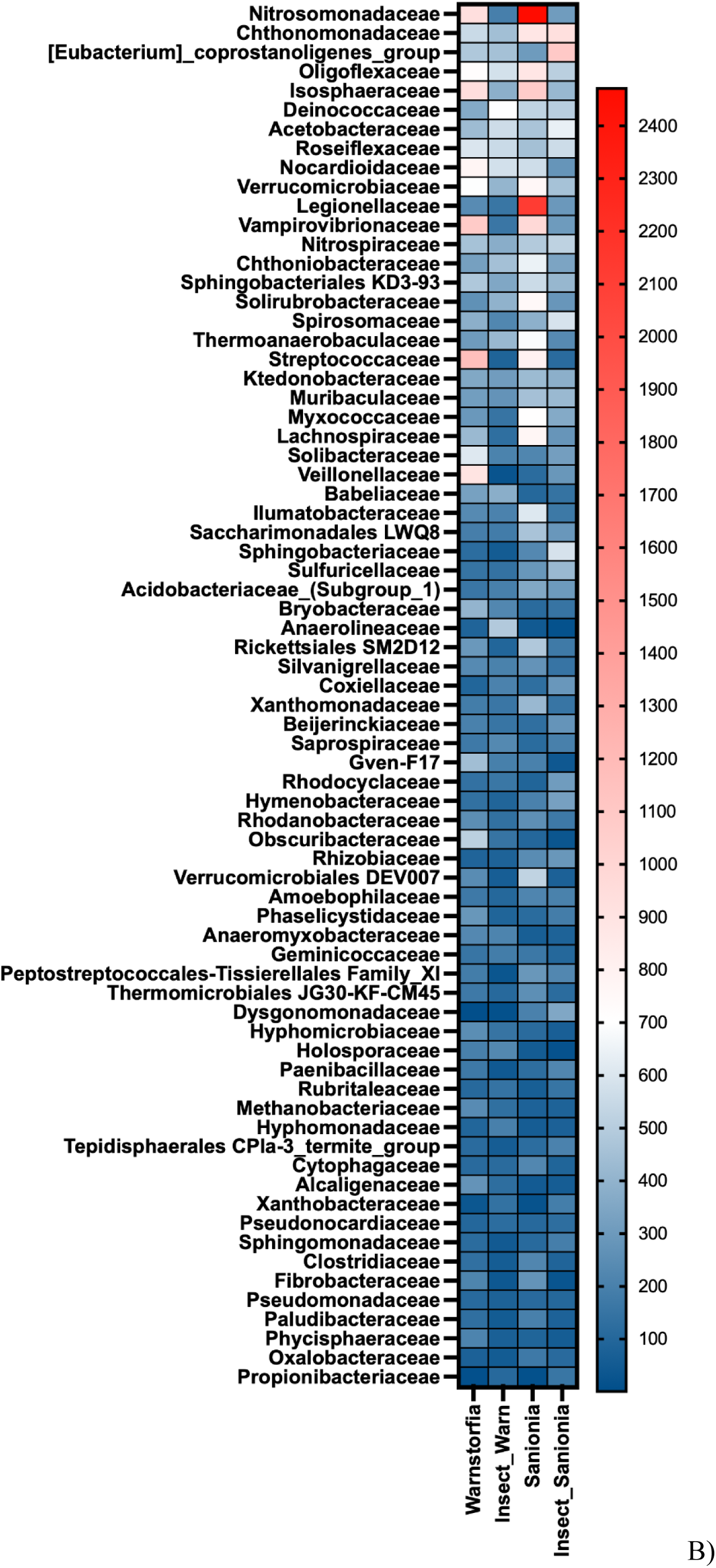
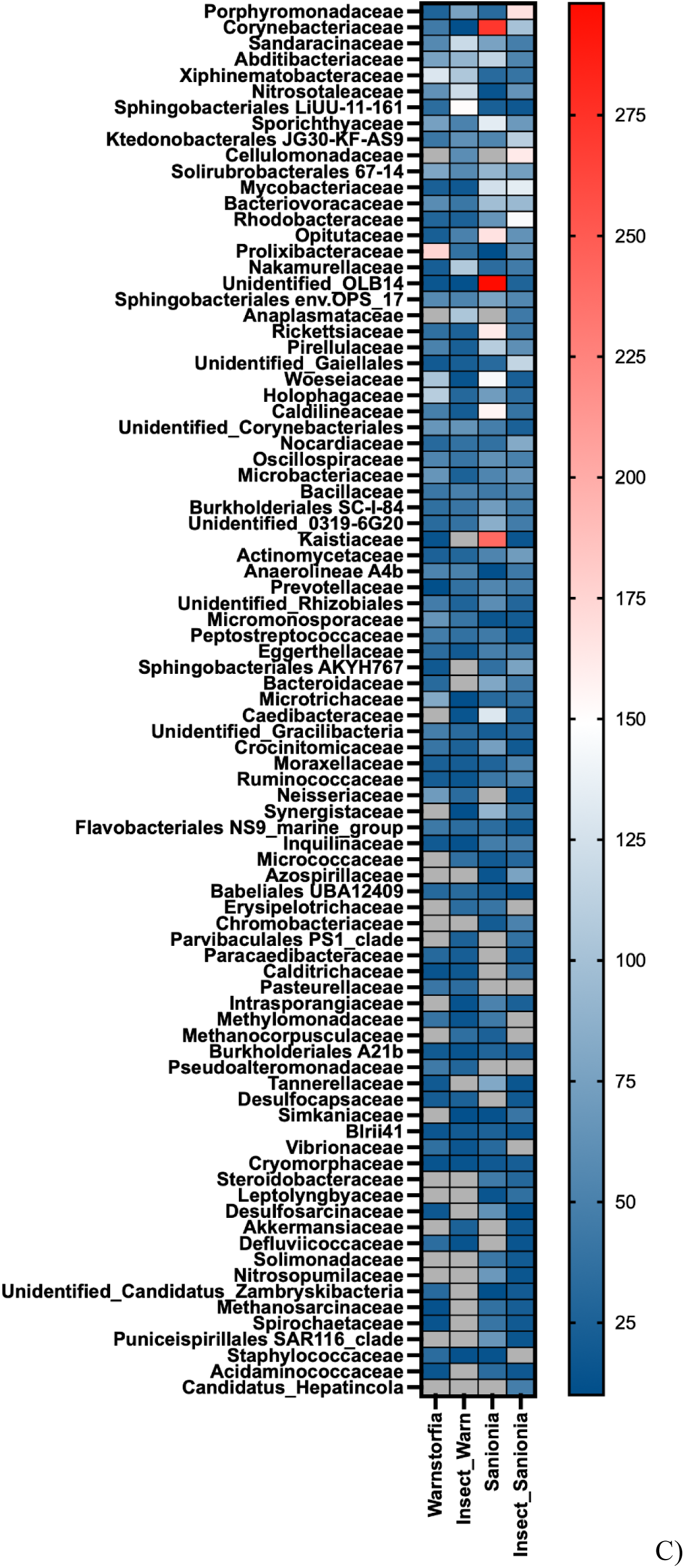
Taxonomic distribution (number of reads) of microbial communities at the family level in different types of samples (Warnstorfia – *Warnstorfia fontinaliopsis moss,* Sanionia *– Sanionia georgicouncinata moss,* InsectWarn – *Belgica antarctica from* the *Warnstorfia fontinaliopsis,* InsectSanionia – *Belgica antarctica* from the *Sanionia georgicouncinata*): A) average abundance < 21236 reads, B) average abundance < 2470 reads , C) average abundance < 297.15 reads

## Notes

### Competing Interest Statement

The authors have declared no competing interest.

## References

1. Adam MAC, Cailleau G, Junier P, Benrey B. Host diet and species interact to shape the bacterial and fungal microbiome in the regurgitant of four *Spodoptera* species. Microb Ecol 2025;88 (1):78. 10.1007/s00248-025-02582-5.

2. Allegrucci G, Carchini G, Convey P, Sbordoni V. Evolutionary geographic relationships among orthocladine chironomid midges from maritime Antarctic and sub-Antarctic islands: Phylogeography in antarctic midges. Biological Journal of the Linnean Society 2012;106 (2):258–274. 10.1111/j.1095-8312.2012.01864.x.

3. Atchley WR, Davis BL. Chromosomal variability in the Antarctic insect, *Belgica antarctica* (Diptera: Chironomidae). Annals of the Entomological Society of America 1979;72 (2):246–252. 10.1093/aesa/72.2.246.

4. Baldo L, Ayoub NA, Hayashi CY, Russell JA, Stahlhut JK, Werren JH. Insight into the routes of *Wolbachia* invasion: high levels of horizontal transfer in the spider genus *Agelenopsis* revealed by *Wolbachia* strain and mitochondrial DNA diversity. Molecular Ecology 2008;17 (2):557–569. 10.1111/j.1365-294X.2007.03608.x.

5. Bang C, Dagan T, Deines P, Dubilier N, Duschl WJ, Fraune S, et al. Metaorganisms in extreme environments: do microbes play a role in organismal adaptation? Zoology 2018;127:1–19. 10.1016/j.zool.2018.02.004.

6. Bargagli R. Terrestrial ecosystems of the Antarctic Peninsula and their responses to climate change and anthropogenic impacts. Ukrainian Antarctic Journal 2020;(2):84–97. 10.33275/1727-7485.2.2020.656.

7. Bartlett JC, Convey P, Hayward SAL. Life cycle and phenology of an Antarctic invader: the flightless chironomid midge, *Eretmoptera murphyi*. Polar Biol 2019;42 (1):115–130. 10.1007/s00300-018-2403-5.

8. Baust JG, Lee RE. Population differences in antifreeze/cryoprotectant accumulation patterns in an Antarctic insect. Oikos 1983;40 (1):120–124. 10.2307/3544206.

9. Baust JG, Lee RE. Multiple stress tolerance in an antarctic terrestrial arthropod: *Belgica antarctica*. Cryobiology 1987;24 (2):140–147. 10.1016/0011-2240(87)90016-2.

10. Bokveld A, Nnolim NE, Nwodo UU. *Chryseobacterium aquifrigidense* FANN1 produced detergent-stable metallokeratinase and amino acids through the abasement of chicken feathers. Front Bioeng Biotechnol 2021;9. 10.3389/fbioe.2021.720176.

11. Bolyen E, Rideout JR, Dillon MR, Bokulich NA, Abnet CC, Al-Ghalith GA, et al. Reproducible, interactive, scalable and extensible microbiome data science using QIIME 2. Nat Biotechnol 2019;37 (8):852–857. 10.1038/s41587-019-0209-9.

12. Braglia C, Cutajar S, Magagnoli S, Asciano D, Burgio G, Di Gioia D, et al. The ground beetle *Poecilus* (Carabidae) gut microbiome and its functionality. Microb Ecol 2025;88 (1):83. 10.1007/s00248-025-02579-0.

13. Brayley ODM, McCready K, Liu S, Convey P, Chen Y, Ullah S, et al. The microbiome of an invasive Antarctic insect, Eretmoptera murphyi (Diptera: Chironomidae), and its potential role in nutrient cycling. 2025a; 10.21203/rs.3.rs-7744438/v1.

14. Brayley O, Hayward S, McCready K, Liu S, Chen Y, Ullah S, et al. Absence of *Wolbachia* in the sub-Antarctic midge, *Eretmoptera murphyi* (Diptera: Chironomidae). Antarctic Science 2025b;37 (4):332–337. 10.1017/S0954102025000197.

15. Brumin M, Kontsedalov S, Ghanim M. *Rickettsia* influences thermotolerance in the whitefly *Bemisia tabaci* B biotype. Insect Science 2011;18 (1):57–66. 10.1111/j.1744-7917.2010.01396.x.

16. Cai B, Han Y, Liu B, Ren Y, Jiang S. Isolation and characterization of an atrazine-degrading bacterium from industrial wastewater in China. Lett Appl Microbiol 2003;36 (5):272–276. 10.1046/j.1472-765X.2003.01307.x.

17. Caporaso JG, Lauber CL, Walters WA, Berg-Lyons D, Lozupone CA, Turnbaugh PJ, et al. Global patterns of 16S rRNA diversity at a depth of millions of sequences per sample. Proc Natl Acad Sci USA 2011;108 (supplement_1):4516–4522. 10.1073/pnas.1000080107.

18. Castro MF, Neves JCL, Francelino MR, Schaefer CEGR, Oliveira TS. Seabirds enrich Antarctic soil with trace metals in organic fractions. Science of The Total Environment 2021;785:147271. 10.1016/j.scitotenv.2021.147271.

19. Chabanol E, Gendrin M. Insects and microbes: best friends from the nursery. Current Opinion in Insect Science 2024;66:101270. 10.1016/j.cois.2024.101270.

20. Charlesworth J, Weinert LA, Araujo EV, Welch JJ. *Wolbachia* , *Cardinium* and climate: an analysis of global data. Biol Lett 2019;15 (8):20190273. 10.1098/rsbl.2019.0273.

21. Chigira A, Miura K. Detection of ‘*Candidatus* Cardinium’ bacteria from the haploid host *Brevipalpus californicus* (Acari: Tenuipalpidae) and effect on the host. Exp Appl Acarol 2005;37 (1–2):107–116. 10.1007/s10493-005-0592-4.

22. Chown SL, Clarke A, Fraser CI, Cary SC, Moon KL, McGeoch MA. The changing form of Antarctic biodiversity. Nature 2015;522 (7557):431–438. 10.1038/nature14505.

23. Convey P, Block W. Antarctic diptera: Ecology, physiology and distribution. EJE 2013;93 (1):1–13.

24. Convey P, Biersma EM. Antarctic Ecosystems. Encyclopedia of Biodiversity 2024;133–148. 10.1016/B978-0-12-822562-2.00058-X.

25. Convey P, Chown SL, Clarke A, Barnes DKA, Bokhorst S, Cummings V, et al. The spatial structure of Antarctic biodiversity. Ecological Monographs 2014;84 (2):203–244. 10.1890/12-2216.1.

26. Convey P, Peck LS. Antarctic environmental change and biological responses. Sci Adv 2019;5 (11):eaaz0888. 10.1126/sciadv.aaz0888.

27. Cranston PS. *Eretmoptera murphy* Schaeffer (Diptera: Chironomidae), an apparently parthenogenetic Antarctic midge. British Antarctic Survey Bulletin 1985;66:35–45.

28. Doremus MR, Kelly SE, Hunter MS. Exposure to opposing temperature extremes causes comparable effects on *Cardinium* density but contrasting effects on *Cardinium*-induced cytoplasmic incompatibility. PLoS Pathog 2019;15 (8):e1008022. 10.1371/journal.ppat.1008022.

29. Doremus MR, Stouthamer CM, Kelly SE, Schmitz-Esser S, Hunter MS. Quality over quantity: unraveling the contributions to cytoplasmic incompatibility caused by two coinfecting *Cardinium* symbionts. Heredity 2022;128 (3):187–195. 10.1038/s41437-022-00507-3.

30. dos Santos Morais G, Vieira TB, Santos GS, Dolatto RG, Cestari MM, Grassi MT, et al. Genotoxic, metabolic, and biological responses of *Chironomus sancticaroli* Strixino & Strixino, 1981 (Diptera: Chironomidae) after exposure to BBP. Science of The Total Environment 2020;715:136937. 10.1016/j.scitotenv.2020.136937.

31. Douglas AE. Multiorganismal insects: Diversity and function of resident microorganisms. Annu Rev Entomol 2015;60 (1):17–34. 10.1146/annurev-ento-010814-020822.

32. Duron O, Bouchon D, Boutin S, Bellamy L, Zhou L, Engelstädter J, et al. The diversity of reproductive parasites among arthropods: *Wolbachia* do not walk alone. BMC Biol 2008;6 (1):27. 10.1186/1741-7007-6-27.

33. Edgar RC. UPARSE: Highly accurate OTU sequences from microbial amplicon reads. Nat Methods 2013;10 (10):996–998. 10.1038/nmeth.2604.

34. Ferree PM, Avery A, Azpurua J, Wilkes T, Werren JH. A Bacterium targets maternally inherited centrosomes to kill males in *Nasonia*. Current Biology 2008;18 (18):1409–1414. 10.1016/j.cub.2008.07.093.

35. Fraser CI, Morrison AK, Hogg AM, Macaya EC, Van Sebille E, Ryan PG, et al. Antarctica’s ecological isolation will be broken by storm-driven dispersal and warming. Nature Clim Change 2018;8 (8):704–708. 10.1038/s41558-018-0209-7.

36. Fujii S, Kawai K, Sambongi Y, Wakai S. Species-specific microorganisms in acid-tolerant *Chironomus* larvae reared in a neutral pH range under laboratory conditions: Single dataset analysis. Microbes Environ 2023;38 (6):ME23029. 10.1264/jsme2.ME23029.

37. Gantz JD, Philip BN, Teets NM, Kawarasaki Y, Potts LJ, Spacht DE, et al. Brief exposure to a diverse range of environmental stress enhances stress tolerance in the polyextremophilic Antarctic midge, Belgica antarctica. 2020;2020.01.01.887414. 10.1101/2020.01.01.887414.

38. Garrido-Bautista J, Norte AC, Moreno-Rueda G, Nadal-Jiménez P. Ecological determinants of prevalence of the male-killing bacterium *Arsenophonus nasoniae*. Journal of Invertebrate Pathology 2024;203:108073. 10.1016/j.jip.2024.108073.

39. Gasparich GE. Spiroplasmas and phytoplasmas: Microbes associated with plant hosts. Biologicals 2010;38 (2):193–203. 10.1016/j.biologicals.2009.11.007.

40. George F, Mahieux S, Daniel C, Titécat M, Beauval N, Houcke I, et al. Assessment of Pb(II), Cd(II), and Al(III) removal capacity of bacteria from food and gut ecological niches: Insights into biodiversity to limit intestinal biodisponibility of toxic metals. Microorganisms 2021;9 (2):456. 10.3390/microorganisms9020456.

41. Gohl P, LeMoine CMR, Cassone BJ. Diet and ontogeny drastically alter the larval microbiome of the invertebrate model *Galleria mellonella*. Can J Microbiol 2022;68 (9):594–604. 10.1139/cjm-2022-0058.

42. Grzesiak J, Kaczyńska A, Gawor J, Żuchniewicz K, Aleksandrzak-Piekarczyk T, Gromadka R, et al. A smelly business: Microbiology of Adélie penguin guano (Point Thomas rookery, Antarctica). Science of The Total Environment 2020;714:136714. 10.1016/j.scitotenv.2020.136714.

43. Haghshenas-Gorgabi N, Poorjavd N, Khajehali J, Wybouw N. *Cardinium* symbionts are pervasive in Iranian populations of the spider mite *Panonychus ulmi* despite inducing an infection cost and no demonstrable reproductive phenotypes when *Wolbachia* is a symbiotic partner. Exp Appl Acarol 2023;91 (3):369–380. 10.1007/s10493-023-00840-0.

44. Haider K, Abbas D, Galian J, Ghafar MA, Kabir K, Ijaz M, et al. The multifaceted roles of gut microbiota in insect physiology, metabolism, and environmental adaptation: implications for pest management strategies. World J Microbiol Biotechnol 2025;41 (3):75. 10.1007/s11274-025-04288-9.

45. Halpern M, Senderovich Y. Chironomid microbiome. Microb Ecol 2015;70 (1):1–8. 10.1007/s00248-014-0536-9.

46. Holmes CJ, Jennings EC, Gantz JD, Spacht D, Spangler AA, Denlinger DL, et al. The Antarctic mite, *Alaskozetes antarcticus*, shares bacterial microbiome community membership but not abundance between adults and tritonymphs. Polar Biol 2019;42 (11):2075–2085. 10.1007/s00300-019-02582-5.

47. Hubert J, Nesvorna M, Klimov PB, Erban T, Sopko B, Dowd SE, et al. Interactions of the intracellular bacterium *Cardinium* with its host, the house dust mite *Dermatophagoides farinae* , based on gene expression data. mSystems 2021;6 (6):e00916–21. 10.1128/mSystems.00916-21.

48. Hughes K, Chwedorzewska K, Molina-Montenegro M, Pertierra LR. Terrestrial non-native species in Antarctica: introduction, impact and management response. 2023; 10.48361/QBTD-QT57.

49. Jõesaar M, Viggor S, Heinaru E, Naanuri E, Mehike M, Leito I, et al. Strategy of *Pseudomonas pseudoalcaligenes* C70 for effective degradation of phenol and salicylate. PLoS ONE 2017;12 (3):e0173180. 10.1371/journal.pone.0173180.

50. Jolliffe IT. Principal component analysis. 2002; 10.1007/b98835.

51. Jung H, Lee D, Lee S, Kong HJ, Park J, Seo Y-S. Comparative genomic analysis of *Chryseobacterium* species: deep insights into plant-growth-promoting and halotolerant capacities. Microbial Genomics 2023;9 (10):001108. 10.1099/mgen.0.001108.

52. Kakizawa S, Hosokawa T, Oguchi K, Miyakoshi K, Fukatsu T. *Spiroplasma* as facultative bacterial symbionts of stinkbugs. Front Microbiol 2022;13. 10.3389/fmicb.2022.1044771.

53. Kaur R, Shropshire JD, Cross KL, Leigh B, Mansueto AJ, Stewart V, et al. Living in the endosymbiotic world of *Wolbachia*: A centennial review. Cell Host & Microbe 2021;29 (6):879–893. 10.1016/j.chom.2021.03.006.

54. Kim O-S, Chae N, Lim HS, Cho A, Kim JH, Hong SG, et al. Bacterial diversity in ornithogenic soils compared to mineral soils on King George Island, Antarctica. J Microbiol 2012;50 (6):1081–1085. 10.1007/s12275-012-2655-7.

55. Kolasa M, Krovi RS, Plewa R, Jaworski T, Kadej M, Smolis A, et al. Host trees partially explain the complex bacterial communities of two threatened saproxylic beetles. Insect Molecular Biology 2025;34 (2):311–321. 10.1111/imb.12973.

56. Kolodny O, Callahan BJ, Douglas AE. The role of the microbiome in host evolution. Phil Trans R Soc B 2020;375 (1808):20190588. 10.1098/rstb.2019.0588.

57. Konai M, Clark EA, Camp M, Koeh AL, Whitcomb RF. Temperature ranges, growth optima, and growth rates of *Spiroplasma* (Spiroplasmataceae , class Mollicutes) species. Current Microbiology 1996;32 (6):314–319. 10.1007/s002849900056.

58. Korczak-Abshire M, Hinke JT, Milinevsky G, Juáres MA, Watters GM. Coastal regions of the northern Antarctic Peninsula are key for gentoo populations. Biol Lett 2021;17 (1):20200708. 10.1098/rsbl.2020.0708.

59. Kovalenko P, Trokhymets V, Parnikoza I, Protsenko Y, Salganskiy O, Dzhulai A, et al. Current status of *Belgica antarctica* Jacobs, 1900 (Diptera: Chironomidae) distribution by the data of Ukrainian Antarctic Expeditions. Ukrainian Antarctic Journal 2021;(2):76–93. 10.33275/1727-7485.2.2021.679.

60. Kozeretska I, Serga S, Kovalenko P, Gorobchyshyn V, Convey P. *Belgica antarctica* (Diptera: Chironomidae): A natural model organism for extreme environments. Insect Science 2022;29 (1):2–20. 10.1111/1744-7917.12925.

61. Kruskal WH, Wallis WA. Use of Ranks in One-criterion variance analysis. Journal of the American Statistical Association 1952;47 (260):583–621. 10.1080/01621459.1952.10483441.

62. Laviad-Shitrit S, Sharaby Y, Sela R, Thorat L, Nath BB, Halpern M. Copper and chromium exposure affect chironomid larval microbiota composition. Science of The Total Environment 2021;771:145330. 10.1016/j.scitotenv.2021.145330.

63. Legendre P, Legendre L. Numerical Ecology. 2012. Available from: https://shop.elsevier.com/books/numerical-ecology/legendre/978-0-444-53868-0

64. Leo C, Nardi F, Cucini C, Frati F, Convey P, Weedon JT, et al. Evidence for strong environmental control on bacterial microbiomes of Antarctic springtails. Sci Rep 2021;11 (1):2973. 10.1038/s41598-021-82379-x.

65. Liu B, Ren Y-S, Su C-Y, Abe Y, Zhu D-H. Pangenomic analysis of *Wolbachia* provides insight into the evolution of host adaptation and cytoplasmic incompatibility factor genes. Front Microbiol 2023;14:1084839. 10.3389/fmicb.2023.1084839.

66. Lo W-S, Ku C, Chen L-L, Chang T-H, Kuo C-H. Comparison of metabolic capacities and inference of gene content evolution in mosquito-associated *Spiroplasma diminutum* and *S. taiwanense*. Genome Biology and Evolution 2013;5 (8):1512–1523. 10.1093/gbe/evt108.

67. Lynch HJ, Naveen R, Trathan PN, Fagan WF. Spatially integrated assessment reveals widespread changes in penguin populations on the Antarctic Peninsula. Ecology 2012;93 (6):1367–1377. 10.1890/11-1588.1.

68. Ma W -J., Schwander T. Patterns and mechanisms in instances of endosymbiont-induced parthenogenesis. J of Evolutionary Biology 2017;30 (5):868–888. 10.1111/jeb.13069.

69. Magoč T, Salzberg SL. FLASH: fast length adjustment of short reads to improve genome assemblies. Bioinformatics 2011;27 (21):2957–2963. 10.1093/bioinformatics/btr507.

70. Maistrenko OM, Serga SV, Kovalenko PA, Kozeretska IA. Bacteria associated with the Antarctic endemic insect *Belgica antarctica* Jacobs (Diptera Chironomidae). Cytol Genet 2023;57 (3):207–212. 10.3103/S0095452723030064.

71. Massey JH, Newton ILG. Diversity and function of arthropod endosymbiont toxins. Trends in Microbiology 2022;30 (2):185–198. 10.1016/j.tim.2021.06.008.

72. McFall-Ngai M, Hadfield MG, Bosch TCG, Carey HV, Domazet-Lošo T, Douglas AE, et al. Animals in a bacterial world, a new imperative for the life sciences. Proc Natl Acad Sci USA 2013;110 (9):3229–3236. 10.1073/pnas.1218525110.

73. McQueen JP, Gattoni K, Gendron EMS, Schmidt SK, Sommers P, Porazinska DL. External and internal microbiomes of Antarctic nematodes are distinct, but more similar to each other than the surrounding environment. Journal of Nematology 2023;55 (1):20230004. 10.2478/jofnem-2023-0004.

74. McQueen JP, Gattoni K, Gendron EMS, Schmidt SK, Sommers P, Porazinska DL. Host identity is the dominant factor in the assembly of nematode and tardigrade gut microbiomes in Antarctic Dry Valley streams. Sci Rep 2022;12 (1):20118. 10.1038/s41598-022-24206-5.

75. Mioduchowska M, Konecka E, Gołdyn B, Pinceel T, Brendonck L, Lukić D, et al. Playing peekaboo with a master manipulator: metagenetic detection and phylogenetic analysis of *Wolbachia* supergroups in freshwater invertebrates. IJMS 2023;24 (11):9400. 10.3390/ijms24119400.

76. Mugo-Kamiri L, Querejeta M, Raymond B, Herniou EA. The effect of diet composition on the diversity of active gut bacteria and on the growth of *Spodoptera exigua* (Lepidoptera: Noctuidae). J Insect Sci 2024;24 (2):13. 10.1093/jisesa/ieae031.

77. Ngando FJ, Tang H, Zhang X, Zhang X, Yang F, Shang Y, et al. Effects of feeding sources and different temperature changes on the gut microbiome structure of *Chrysomya megacephala* (Diptera: Calliphoridae). Insects 2025;16 (3):283. 10.3390/insects16030283.

78. Nowak, K. H., Hartop, E., Prus-Frankowska, M., Buczek, M., Kolasa, M. R., Roslin, T., Ovaskainen, O., & Łukasik, P. (2025). What lurks in the dark? An innovative framework for studying diverse wild insect microbiota. Microbiome, 13(1), 186. 10.1186/s40168-025-02169-9

79. Oksanen J, Simpson GL, Blanchet FG, Kindt R, Legendre P, Minchin PR, et al. Vegan: Community Ecology package. 2025;2.7–2. 10.32614/CRAN.package.vegan.

80. Parnikoza I, Berezkina A, Moiseyenko Y, Malanchuk V, Kunakh V. Complex survey of the Argentine Islands and Galindez Island (Maritime Antarctic) as a research area for studying the dynamics of terrestrial vegetation. Ukrainian Antarctic Journal 2018;(1(17)):73–101. 10.33275/1727-7485.1(17).2018.34.

81. Pascual J, García-López M, González I, Genilloud O. *Luteolibacter gellanilyticus* sp. nov., a gellan-gum-degrading bacterium of the phylum *Verrucomicrobia* isolated from miniaturized diffusion chambers. International Journal of Systematic and Evolutionary Microbiology 2017;67 (10):3951–3959. 10.1099/ijsem.0.002227.

82. Potts LJ, Gantz JD, Kawarasaki Y, Philip BN, Gonthier DJ, Law AD, et al. Environmental factors influencing fine-scale distribution of Antarctica’s only endemic insect. Oecologia 2020;194 (4):529–539. 10.1007/s00442-020-04714-9.

83. Prekrasna-Kviatkovska Y, Parnikoza I, Yerkhova A, Stelmakh O, Pavlovska M, Dzyndra M, et al. From acidophilic to ornithogenic: microbial community dynamics in moss banks altered by gentoo penguins. Front Microbiol 2024;15:1362975. 10.3389/fmicb.2024.1362975.

84. Prieto-Fernández F, Lambert S, Kujala K. Assessment of microbial communities from cold mine environments and subsequent enrichment, isolation and characterization of putative antimony- or copper-metabolizing microorganisms. Front Microbiol 2024;15. 10.3389/fmicb.2024.1386120.

85. Protsenko Y, Protsenko O, Kovalenko P, Parnikoza I, Puhovkin A, Svetlichny L, et al. New Collembola occurrence records from the western Antarctic Peninsula. Ukrainian Antarctic Journal 2025;23 (1):64–89. 10.33275/1727-7485.1.2025.744.

86. Quast C, Pruesse E, Yilmaz P, Gerken J, Schweer T, Yarza P, et al. The SILVA ribosomal RNA gene database project: improved data processing and web-based tools. Nucleic Acids Research 2012;41 (D1):D590–D596. 10.1093/nar/gks1219.

87. Raychoudhury R, Baldo L, Oliveira DCSG, Werren JH. Modes of acquisition of *Wolbachia*: Horizontal transfer, hybrid introgression, and codivergence in the *Nasonia* species complex. Evolution 2009;63 (1):165–183. 10.1111/j.1558-5646.2008.00533.x.

88. Rodrigues J, Lefoulon E, Gavotte L, Perillat-Sanguinet M, Makepeace B, Martin C, et al. *Wolbachia* springs eternal: symbiosis in Collembola is associated with host ecology. R Soc Open Sci 2023;10 (5):230288. 10.1098/rsos.230288.

89. Rosenberg E. The family *Chitinophagaceae*. The Prokaryotes 2014;493–495. 10.1007/978-3-642-38954-2_137.

90. Rouf MA, Rigney MM. Bacterial florae in larvae of the lake fly *Chironomus plumosus*. Applied and Environmental Microbiology 1993;59 (4):1236–1241. 10.1128/aem.59.4.1236-1241.1993.

91. Sela R, Halpern M. The Chironomid microbiome plays a role in protecting its host from toxicants. Front Ecol Evol 2022;10:796830. 10.3389/fevo.2022.796830.

92. Sela R, Laviad-Shitrit S, Thorat L, Nath BB, Halpern M. *Chironomus ramosus* larval microbiome composition provides evidence for the presence of detoxifying enzymes. Microorganisms 2021;9 (8):1571. 10.3390/microorganisms9081571.

93. Senderovich Y, Halpern M. Bacterial community composition associated with chironomid egg masses. J Insect Sci 2012;12 (1):149. 10.1673/031.012.14901.

94. Senderovich Y, Halpern M. The protective role of endogenous bacterial communities in chironomid egg masses and larvae. ISME J 2013;7 (11):2147–2158. 10.1038/ismej.2013.100.

95. Serga S, Kovalenko PA, Maistrenko OM, Deconninck G, Shevchenko O, Iakovenko N, et al. *Wolbachia* in Antarctic terrestrial invertebrates: Absent or undiscovered? Environ Microbiol Rep 2024;16 (6):e70040. 10.1111/1758-2229.70040.

96. Serga SV, Maistrenko OM, Matiytsiv NP, Vaiserman AM, Kozeretska IA. Effects of *Wolbachia* infection on fitness-related traits in *Drosophila melanogaster*. Symbiosis 2021;83 (2):163–172. 10.1007/s13199-020-00743-3.

97. Shapiro SS, Wilk MB. An analysis of variance test for normality (complete samples). Biometrika 1965;52 (3–4):591–611. 10.1093/biomet/52.3-4.591.

98. Siegert M, Atkinson A, Banwell A, Brandon M, Convey P, Davies B, et al. The Antarctic Peninsula under a 1.5°C global warming scenario. Front Environ Sci 2019;7:102. 10.3389/fenvs.2019.00102.

99. Singh A, Misser S, Allam M, Chan W-Y, Ismail A, Munhenga G, et al. The effect of larval exposure to heavy metals on the gut microbiota composition of adult *Anopheles arabiensis* (Diptera: Culicidae). Tropical Medicine and Infectious Disease 2024;9 (10):249. 10.3390/tropicalmed9100249.

100. Soares TA, Souza-Kasprzyk J, De Assis Guilherme Padilha J, Convey P, Costa ES, Torres JPM. Ornithogenic mercury input to soils of Admiralty Bay, King George Island, Antarctica. Polar Biol 2024;47 (9):891–901. 10.1007/s00300-023-03162-4.

101. Socolovschi C, Gaudart J, Bitam I, Huynh TP, Raoult D, Parola P. Why Are there so few *Rickettsia conorii conorii*-infected *Rhipicephalus sanguineus* ticks in the wild? PLOS Neglected Tropical Diseases 2012;6 (6):e1697. 10.1371/journal.pntd.0001697.

102. Sugg P, Edwards JS, Baust J. Phenology and life history of *Belgica antarctica* , an Antarctic midge (Diptera: Chironomidae). Ecological Entomology 1983;8 (1):105–113. 10.1111/j.1365-2311.1983.tb00487.x.

103. Turner J, Barrand NE, Bracegirdle TJ, Convey P, Hodgson DA, Jarvis M, et al. Antarctic climate change and the environment: an update. Polar Record 2014;50 (3):237–259. 10.1017/S0032247413000296.

104. Usher MB, Edwards M. A dipteran from south of the Antarctic Circle: *Belgica antarctica* (Chironomidae) with a description of its larva. Biological Journal of the Linnean Society 1984;23 (1):19–31. 10.1111/j.1095-8312.1984.tb00803.x.

105. van den Ende C, White LT, van Welzen PC. The existence and break-up of the Antarctic land bridge as indicated by both amphi-Pacific distributions and tectonics. Gondwana Research 2017;44:219–227. 10.1016/j.gr.2016.12.006.

106. Vega GC, Convey P, Hughes KA, Olalla-Tárraga MÁ. Humans and wind, shaping Antarctic soil arthropod biodiversity. Insect Conserv Diversity 2020;13 (1):63–76. 10.1111/icad.12375.

107. Wasimuddin, Schlaeppi K, Ronchi F, Leib SL, Erb M, Ramette A. Evaluation of primer pairs for microbiome profiling from soils to humans within the One Health framework. Molecular Ecology Resources 2020;20 (6):1558–1571. 10.1111/1755-0998.13215.

108. Weinert LA, Araujo-Jnr EV, Ahmed MZ, Welch JJ. The incidence of bacterial endosymbionts in terrestrial arthropods. Proc R Soc B 2015;282 (1807):20150249. 10.1098/rspb.2015.0249.

109. Wernegreen JJ. Mutualism meltdown in insects: bacteria constrain thermal adaptation. Current Opinion in Microbiology 2012;15 (3):255–262. 10.1016/j.mib.2012.02.001.

110. Werren JH, Baldo L, Clark ME. *Wolbachia*: Master manipulators of invertebrate biology. Nat Rev Microbiol 2008;6 (10):741–751. 10.1038/nrmicro1969.

111. Werren JH, Skinner SW, Huger AM. Male-killing bacteria in a parasitic wasp. Science 1986;231 (4741):990–992. 10.1126/science.3945814.

112. Wilde J, Slack E, Foster KR. Host control of the microbiome: Mechanisms, evolution, and disease. Science 2024;385 (6706):eadi3338. 10.1126/science.adi3338.

113. Williams PA, Sayers JR. The evolution of pathways for aromatic hydrocarbon oxidation in *Pseudomonas*. Biodegradation 1994;5 (3–4):195–217. 10.1007/BF00696460.

